# Multimodal fusion analysis of functional, cerebrovascular and structural neuroimaging in healthy ageing subjects

**DOI:** 10.1101/2021.12.22.473894

**Authors:** Xulin Liu, Lorraine K Tyler, Cam-CAN, James B Rowe, Kamen A Tsvetanov

## Abstract

Brain ageing is a complex process which requires a multimodal approach. Neuroimaging can provide insights into brain morphology, functional organization and vascular dynamics. However, most neuroimaging studies of ageing have focused on each imaging modality separately, limiting the understanding of interrelations between processes identified by different modalities and their relevance to cognitive decline in ageing. Here, we used a data-driven multimodal approach, linked independent component analysis (ICA), to jointly analyze magnetic resonance imaging of grey matter volume, cerebrovascular, and functional network topographies in relation to measures of fluid intelligence. Neuroimaging and cognitive data from the Cambridge Centre for Ageing and Neuroscience study were used, with healthy participants aged 18 to 88 years (main dataset *n* = 215; secondary dataset *n* = 433). Using linked ICA, functional network activities were characterized in independent components but not captured in the same component as structural and cerebrovascular patterns. Split-sample (*n* = 108/107) and out-of-sample (*n* = 433) validation analyses using linked ICA were also performed. Global grey matter volume with regional cerebrovascular changes and the right frontoparietal network activity were correlated with age-related and individual differences in fluid intelligence. This study presents the insights from linked ICA to bring together measurements from multiple imaging modalities, with independent and additive information. We propose that integrating multiple neuroimaging modalities allows better characterization of brain pattern variability and changes associated with healthy ageing.

## 1. INTRODUCTION

Increasing life expectancy is leading to rapid ageing of the worldwide population (Beard et al., 2016). The quality of these extra years of life heavily depends on good health, including the maintenance of good cognitive function across the lifespan (Beard et al., 2016; Sahakian, 2014). There is a pressing need to better understand the neurobiology of cognitive function associated with ageing. Neuroimaging studies show age-related changes in brain morphology, functional networks, and vascular dynamics (Kennedy & Raz, 2015). However, these effects are usually studied separately, whereas their integration could explain how these components influence cognitive ageing (K. A. Tsvetanov, Henson, & Rowe, 2021).

Brain atrophy is one of the most commonly studied features of ageing (Grajauskas et al., 2019; Pini et al., 2016; Romanowski & Wilkinson, 2011). Atrophy refers to the loss of brain tissue, which mainly comprises of grey matter constituted by the cell bodies of neurons and glial cells (Azevedo et al., 2009). The concentration and volume of grey matter can be estimated from structural magnetic resonance imaging (MRI) and generally decrease with age (Grajauskas et al., 2019; Kennedy & Raz, 2015). Nevertheless, these changes are not uniform across the brain regions and patterns of brain ageing vary among individuals and the trajectories can be influenced by environmental, genetic and other neurobiological factors (Bethlehem et al., 2022; Lemaitre et al., 2012; Pini et al., 2016). In fact, atrophy on its own does not fully explain cognitive performance (Boekel et al., 2015) and is insufficient for understanding ageing and neurodegenerative syndromes with heterogenous clinical features (Grajauskas et al., 2019; Murley et al., 2020; Perry et al., 2017; K. A. Tsvetanov, Gazzina, et al., 2021; K. A. Tsvetanov et al., 2016). Instead, we propose that brain ageing is multifactorial, reflecting complex processes which require multivariate techniques to elucidate (Doan, Engvig, Persson, et al., 2017; Doan, Engvig, Zaske, et al., 2017; Douaud et al., 2014; Groves, Beckmann, Smith, & Woolrich, 2011; Murley et al., 2020). Neuroimaging is a key contributor to this approach, from its quantification of brain morphology, functional networks and vascular dynamics.

Brain functional networks are commonly studied using functional magnetic resonance imaging (fMRI), which measures neural activity indirectly via changes in the blood oxygen level-dependent (BOLD) signal (Chen & Glover, 2015; Grady, 2012; Rosen & Savoy, 2012). Cognitive function is dependent on intrinsic interactions within large-scale functional brain networks as well as extrinsic interactions between such functional brain networks (Fox et al., 2005; Kelly, Uddin, Biswal, Castellanos, & Milham, 2008). These networks show selective vulnerability to age and neurodegeneration (Moguilner et al., 2020; K. A. Tsvetanov, Gazzina, et al., 2021). Task-free fMRI, also known as resting-state fMRI (rs-fMRI), can be used to characterize intrinsic and extrinsic connectivity of functional networks simultaneously (Cole, Bassett, Power, Braver, & Petersen, 2014; Smith et al., 2009). Spontaneous activity, which can be measured by rs-fMRI, is the most metabolic demanding component of neural activity (Raichle & Mintun, 2006). Moreover, activities in resting-state functional networks, such as the default mode network (DMN), the salience network (SN) and the frontoparietal network (FPN), are associated with a wide range of cognitive functions (e.g., memory, language, attention, visual processes) (Corbetta & Shulman, 2002) and playing an increasingly important role in maintaining good cognition in old age and progression of some neurodegenerative diseases (Bethlehem et al., 2020; Day et al., 2013; Marek & Dosenbach, 2018; Rosazza & Minati, 2011; Tibon et al., 2021; K. A. Tsvetanov, Gazzina, et al., 2021; K. A. Tsvetanov et al., 2016; Zhou & Seeley, 2014).

Resting-state fMRI BOLD signals also reflect the haemodynamic response evoked by neuronal activity and therefore they represent both vascular and neuronal signals (K. A. Tsvetanov, Henson, & Rowe, 2021). Age differences in BOLD signal could be confounded by differences in cerebrovascular function associated with non-neuronal physiological factors (e.g., medications, time of day, or level of wakefulness). Nevertheless, cerebrovascular function is also implicated as a major factor in maintaining brain health in ageing and neurodegenerative diseases (Barisano et al., 2022; Fuhrmann et al., 2019; Iadecola, 2017; Kisler, Nelson, Montagne, & Zlokovic, 2017; Sweeney, Kisler, Montagne, Toga, & Zlokovic, 2018; Zimmerman, Rypma, Gratton, & Fabiani, 2021). The mixture of cerebrovascular and neuronal contributions to BOLD signals complicates the interpretation of age differences in BOLD imaging and, thus, our understanding of neurocognitive ageing (K. A. Tsvetanov et al., 2015). Dissociation of vascular and neuronal signals would therefore be particularly meaningful (K. A. Tsvetanov, Henson, & Rowe, 2021).

An important aspect of cerebrovascular function is the ability to move blood through a network of cerebral vasculature supplying the brain, which can be assessed by measuring resting cerebral blood flow (CBF) using arterial spin labelling (ASL) (Detre, Wang, Wang, & Rao, 2009; Williams, Detre, Leigh, & Koretsky, 1992). ASL is a noninvasive MRI technique used to quantify cerebral blood perfusion by labelling blood water as it flows throughout the brain. Baseline CBF, which relates to cerebrovascular ageing (K. A. Tsvetanov, Henson, Jones, et al., 2021), is important for maintaining cognitive function, while its non-neuronal contributions to BOLD signals reflect age-related confound (Wu et al., 2021). Another important aspect of cerebrovascular function is the ability to regulate regional blood flow via carbon dioxide-modulated constriction or dilation of cerebral vessels (Willie, Tzeng, Fisher, & Ainslie, 2014). This cerebrovascular reactivity can be assessed using resting state fluctuation amplitude (RSFA) (Kannurpatti & Biswal, 2008), which reflects naturally occurring fluctuations in BOLD signals induced by variations in cardiac and respiratory rhythms at resting state (Birn, Diamond, Smith, & Bandettini, 2006; Shmueli et al., 2007). RSFA is a safe, scalable, and robust alternative to hypercapnia and breath-holding approaches (Kannurpatti & Biswal, 2008; K. A. Tsvetanov, Henson, Jones, et al., 2021). RSFA is sensitive to cardiovascular and cerebrovascular differences in ageing (Garrett, Lindenberger, Hoge, & Gauthier, 2017; K. A. Tsvetanov et al., 2015; K. A. Tsvetanov, Henson, Jones, et al., 2021), intracranial vascular disease (Makedonov, Black, & MacIntosh, 2013; Nair, Raut, & Prabhakaran, 2017; Raut, Nair, Sattin, & Prabhakaran, 2016), neurodegeneration (Makedonov, Chen, Masellis, & MacIntosh, 2016; Peter R. Millar et al., 2020) and cognitive impairment (Millar et al., 2021; Peter R Millar et al., 2020; Kamen A Tsvetanov et al., 2022).

The majority of neuroimaging studies have focused on each imaging modality separately, limiting the understanding of interrelations between modalities and the complex neural mechanisms associated with cognitive change. Unraveling the interactive effects of changes on morphometry, cerebrovascular and functional levels could provide better understanding of the multifactorial neurobiological mechanisms underlying cognitive change in ageing and neurodegeneration. Linked independent component analysis (ICA) is a data-driven analytic method that allows for simultaneous characterization of multimodal imaging modalities while taking into account the covariance across imaging modalities (Groves et al., 2011). In comparison with other commonly used multivariate approaches such as canonical correlation analysis (CCA) and partial least squares (PLS), linked ICA is able to identify patterns of covariance across more than two modalities. By identifying common patterns that are shared by different imaging modalities and identifying independent components that are dominated by single imaging modality, one can more accurately characterize the predictors of the outcomes of interest.

We aimed to integrate structural, functional and cerebrovascular neuroimaging signals to better understand their contribution to cognitive decline in ageing. We focused on fluid intelligence, which includes reasoning and problem-solving abilities (Gottfredson & Deary, 2004) and has been shown consistently to decrease markedly with ageing (Ghisletta, Rabbitt, Lunn, & Lindenberger, 2012; Hartshorne & Germine, 2015; Kievit et al., 2014; T. A. Salthouse, 2009; Timothy A. Salthouse, 2010). Specifically, we tested whether differences in structural, cerebrovascular, and functional network activation patterns have independent or convergent patterns with age, and whether these patterns are correlated with measures of fluid intelligence across the lifespan.

## 2. METHODS

### 2.1 Cohorts and participants

The Cambridge Centre for Ageing and Neuroscience (Cam-CAN) cohort study recruited healthy adults from its local general population in the UK, in three stages (Shafto et al., 2014). In Stage 1, 3000 adults aged 18 and above were recruited for a home interview. In Stage 2, a subset of 700 participants aged 18-88 (100 per age decile) was selected to participate in neuroimaging (e.g., structural MRI and fMRI) and behavioural tests (Shafto et al., 2014). We refer to Stage 2 as CC700 in this paper. In Stage 3, a subset of 280 participants (40 per age decile) was selected to participate in further neuroimaging (e.g., fMRI, ASL) and cognitive examinations across key cognitive domains (Shafto et al., 2014; Taylor et al., 2017), and we refer to Stage 3 as CC280 in this paper. Details of the neuroimaging experiments and cognitive tasks are reported previously (Shafto et al., 2014; Taylor et al., 2017). Fluid intelligence was assessed as a principal cognitive measure due to its strong positive correlations with performance on many other cognitive tests, and sensitivity to age. To assess fluid intelligence in the Cam-CAN study, we used the standard form of the Cattell Culture Fair, Scale 2 Form A (Cattell, 1971; Cattell, Cattell, Institute for, & Ability, 1960). This test contained four subtests with different types of nonverbal “puzzles”: series completion, classification, matrices, and conditions. Each subtest was timed with 3 minutes for the first subtest, 4 minutes for the second, 3 minutes for the third, and 2.5 minutes for the final subtest (participants were not informed about precise timings beforehand) (Shafto et al., 2014). Before each subtest, instructions were read from the manual and participants were given examples. The Cattell test was a pen-and-paper test where the participant chose a response on each trial from multiple choices, and recorded responses on an answer sheet. Correct responses were given a score of 1 for a total maximum score of 46. The total score of Cattell test was interpreted as a measure of fluid intelligence in this study. Ethical approval was obtained from the Cambridge 2 Research Ethics Committee, and written informed consent was given by all participants. Subjects in Cam-CAN Stage 3 (CC280) were analyzed as the main analysis of this study (*n* = 215). Subjects from CC700 that were excluded from the CC280 main analysis formed an independent sample for out-of-sample validation analysis, and we refer to this sample as CC420 (*n* = 433) in this paper.

### 2.2 Image acquisition

Imaging data from Cam-CAN were acquired using a 3T Siemens TIM Trio. A 3D structural MRI was acquired using T1-weighted sequence with generalized autocalibrating partially parallel acquisition with acceleration factor 2; repetition time (TR) = 2250 ms; echo time (TE) = 2.99 ms; inversion time (TI) = 900 ms; flip angle α = 9°; field-of-view (FOV) = 256 X 240 X 192 mm; resolution = 1 mm isotropic; acquisition time of 4 min and 32 s.

For rs-fMRI, echoplanar imaging (EPI) acquired 261 volumes with 32 slices (sequential descending order, slice thickness of 3.7 mm with a slice gap of 20% for whole-brain coverage, TR = 1970 ms; TE = 30 ms; flip angle α = 78°; FOV = 192 mm × 192 mm; resolution = 3 mm × 3 mm × 4.44 mm) during 8 min and 40 s. Participants were instructed to lie still with their eyes closed. The initial six volumes were discarded to allow for T1 equilibration.

An index of cerebrovascular reactivity was estimated using the RSFA (Kannurpatti & Biswal, 2008; K. A. Tsvetanov et al., 2015; K. A. Tsvetanov, Henson, & Rowe, 2021). RSFA was estimated from the resting-state EPI reported above. Subject specific RSFA maps were calculated based on the normalized standard deviation across time for processed rs-fMRI time series data. Details on the acquisition of RSFA are also reported previously (K. A. Tsvetanov, Henson, Jones, et al., 2021).

To assess resting CBF, pulsed ASL was used (PASL, PICORE-Q2T-PASL in axial direction, 2,500 ms repetition time, 13 ms echo time, bandwidth 2,232 Hz/Px, 256 × 256 mm2 field of view, imaging matrix 64 × 64, 10 slices, 8 mm slice thickness, flip angle 90°, 700 ms TI1, TI2 = 1,800 ms, 1,600 ms saturation stop time). The imaging volume was positioned to maintain maximal brain coverage with a 20.9 mm gap between the imaging volume and a labeling slab with 100 mm thickness. There were 90 repetitions giving 45 control-tag pairs (duration 3’52”). A single-shot EPI (M0) equilibrium magnetization scan was acquired.

### 2.3 Image preprocessing

Preprocessing of T1-weighted images used standardized preprocessing consistent with Cam-CAN data processing protocol (Taylor et al., 2017; K. A. Tsvetanov, Henson, Jones, et al., 2021). The Automatic Analysis (Cusack et al., 2014) pipelines implemented in Matlab (MathWorks) were used. The T1 image was initially coregistered to the MNI template, and the T2 image was then coregistered to the T1 image using a rigid-body transformation. The coregistered T1 and T2 images were used in a multichannel segmentation to extract probabilistic maps of six tissue classes: grey matter, white matter, cerebrospinal fluid, bone, soft tissue, and residual noise. The native space grey matter and white matter images were submitted to diffeomorphic registration (DARTEL) (Ashburner, 2007) to create group template images. Each template was normalized to the MNI template using a 12-parameter affine transformation. Images were modulated to correct for individual brain size. Grey matter images were smoothed with an 8 mm full-width at half maximum (FWHM) Gaussian kernel in alignment with the standardized processing protocol of Cam-CAN data (Taylor et al., 2017; K. A. Tsvetanov et al., 2018). Modulated grey matter volume (GMV) was analyzed in linked ICA of this study. For the linked ICA grey matter images were down-sampled to match the resolution of fMRI and perfusion data. A brain mask from Statistical Parametric Mapping 12 (SPM12) (https://www.fil.ion.ucl.ac.uk/spm/software/spm12/) was applied at a threshold of 0.9 (i.e., regions with > 90% probability being within the brain were included).

Resting-state fMRI data were preprocessed using Automatic Analysis (Cusack et al., 2014) calling functions from SPM12 implemented in Matlab (MathWorks). Resting-state fMRI were further processed using whole-brain ICA of single-subject time series denoising (termed subject-ICA), with noise components selected and removed automatically using the ICA-based Automatic Removal of Motion Artifacts toolbox (AROMA) (Pruim, Mennes, Buitelaar, & Beckmann, 2015; Pruim, Mennes, van Rooij, et al., 2015). This was complemented with linear detrending of the fMRI signal, covarying out six realignment parameters, white matter and cerebrospinal fluid signals, their first derivatives, and quadratic terms (Pruim, Mennes, van Rooij, et al., 2015). Global white matter and cerebrospinal fluid signals were estimated for each volume from the mean value of white matter and cerebrospinal fluid masks derived by thresholding SPM tissue probability maps at 0.75. Resting-state fMRI data were head motion corrected, bandpass filtered and spatially smoothed with a 6 mm FWHM Gaussian kernel in accordance with recommendations of the AROMA processing pipeline. As the subsequent analysis method is robust to potential differences in spatial smoothness across modalities, we used modality-specific smoothing kernels.

Pulsed ASL time series were converted to CBF maps using ExploreASL toolbox (H. Mutsaerts et al., 2018). Following rigid-body alignment, the images were coregistered with the T1 volume, normalised with normalization parameters from the T1 stream to warp ASL images into MNI space (K. A. Tsvetanov, Henson, Jones, et al., 2021). Given the ASL data was based on a sequence with lower resolution (i.e., slice thickness of 8 mm), a smoothing kernel size 1.5 times larger than the slice thickness (i.e., 12 mm FWHM Gaussian kernel) was used, consistent with the efficacy of ASL data with heavier smoothing kernels (Wang, Wang, Aguirre, & Detre, 2005). RSFA was smoothed with an 8 mm FWHM Gaussian kernel.

All T1 and EPI image processing came from Release004 of the Cam-CAN pipelines, which included quality-control checks by semi-automated scripts monitored by the Cam-CAN methods team (Taylor et al., 2017). CBF images with artefacts (*n* = 25) based on visual inspection were excluded from analysis.

### 2.4 Image analysis

A summary flow chart of the processing and analysis of imaging modalities is presented in Figure 1.

**Figure 1.**
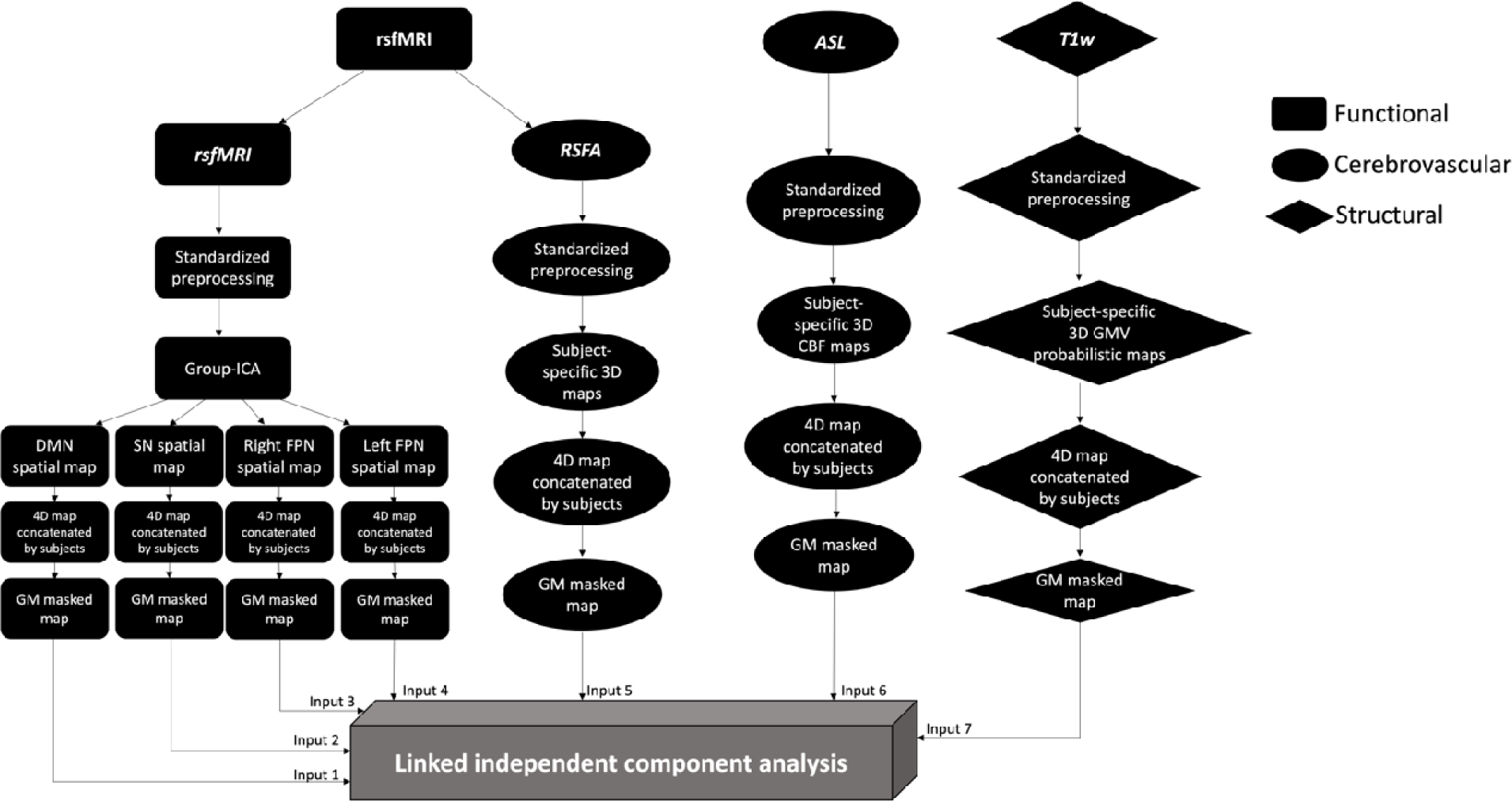
Summary of processing and analysis of the imaging modalities, comprising functional, cerebrovascular and structural measurements. Abbreviations: ASL, arterial spin labelling; *CBF*, cerebral blood flow; *DMN*, default mode network; *FPN*, frontoparietal network; *GMV*, grey matter volume; *ICA*, independent component analysis; *RSFA*, resting state fluctuation amplitude; *rsfMRI*, resting-state functional magnetic resonance imaging; *SN*, salience network; T1w, T1-weighted.

#### 2.4.1 Functional network decomposition using group-ICA

In order to identify functional networks from rs-fMRI and study network spatial patterns, an ICA was performed using the Group-level ICA of fMRI Toolbox to decompose the rs-fMRI (<u>trendscenter.org/software/gift/</u>) (V. D. Calhoun, Adali, Pearlson, & Pekar, 2001). ICA dissociates signals from complex datasets with minimal assumptions (V. Calhoun, 2018), to represent data in a small number of independent components (ICs) which here are spatial maps that describe the temporal and spatial characteristics of underlying signals (V. D. Calhoun et al., 2001; McKeown et al., 1998). The values at each voxel reflect the correlation between the timeseries of the voxel and that of the component. Each component can therefore be interpreted as similar BOLD activity of a functional network at resting-state (Rosazza & Minati, 2011).

The data from participants in CC700 (*n* = 648) were analyzed using group-ICA. This provided a twofold advantage: subjects excluded from the main analysis (CC280) formed an independent second sample (see below in 2.5); and having a larger sample increases the reliability of ICA decomposition results while maximizing statistical power (V. D. Calhoun, Kiehl, & Pearlson, 2008; Erhardt et al., 2011). The number of components used, N = 15, matched a common degree of decomposition previously applied in low-dimensional group-ICA of rs-fMRI (Beckmann, DeLuca, Devlin, & Smith, 2005; Damoiseaux et al., 2006; Smith et al., 2009) and generated network spatial maps that showed a high degree of overlapping with network templates. Low-dimensional group-ICA was used because the purpose was to define each network with a single component, and high-dimensional group-ICA would tend to decompose single network into multiple components. Hundred ICASSO iterations were used to ensure the reliability of estimated ICs (Himberg & Hyvarinen, 2003). Functional networks were identified from components by visualization and validated by spatially matching the components to pre-existing templates (Shirer, Ryali, Rykhlevskaia, Menon, & Greicius, 2012), in accordance with previous methodology used to identify networks from ICs (K. A. Tsvetanov et al., 2016). Four resting-state functional networks were selected to achieve a relatively balanced number of inputs between functional and non-functional imaging measurements. The DMN, SN, right and left FPN were selected, which are higher-order functional networks known to be associated with age and cognitive change including fluid intelligence (Buckner, Andrews-Hanna, & Schacter, 2008; Corbetta & Shulman, 2002; Samu et al., 2017; Snyder, Uddin, & Nomi, 2021; Tibon et al., 2021).

#### 2.4.2 Multimodal fusion using linked ICA

Linked ICA was performed using FLICA of FMRIB (https://fsl.fmrib.ox.ac.uk/fsl/fslwiki/FLICA) (Groves et al., 2011; Smith et al., 2004) implemented in Matlab (MathWorks version 2020b). Linked ICA was run with 7 spatial map inputs: GMV, CBF, RSFA and four maps from three resting-state functional networks (i.e., the DMN, the SN, the right FPN and the left FPN) of those subjects that were included in CC280. We refer to these imaging derived inputs as modalities. Within each modality, images from all subjects were concatenated into a single input image for linked ICA. Linked ICA decomposed this n-by-m matrix of participants-by-voxels into spatial components, with each component being an aggregate of spatial patterns, one for each modality, along with a set of subject loadings, one for each component (for more details see (Groves et al., 2011; Groves et al., 2012)). Each modality spatial pattern is a map of weights that is later converted to pseudo-Z-statistic by accounting the scaling of the variables and the signal-to-noise ratio in that modality. Only modalities with significant contribution (i.e., having weight with Z-score > 3.34, which corresponds to *P* < 0.001) were presented in this study. Linked ICA subject loadings for a given component were shared between all modalities represented in that component and indicated the degree to which that component was presented in any individual subject. Subject loadings were used as inputs to the second-level between-subject regression analysis (see below in 2.6). To ensure that results were not influenced dominantly by non-grey matter regions, a grey matter probability mask from SPM12 was used with a threshold of 0.3. We performed linked ICA using a dimensionality of 40, with 1000 iterations based on recommendation in previous studies (Doan, Engvig, Zaske, et al., 2017; Doan, Kaufmann, et al., 2017; Francx et al., 2016; Groves et al., 2012; Li et al., 2020; Wolfers et al., 2017). To ensure linked ICA fusion patterns were robust to the model order, we also performed the linked ICA using 30 and 50 dimensions. To ensure linked ICA fusion patterns were not biased by multiple functional network inputs, we repeated linked ICA with only one functional network (DMN) input to examine the fusion patterns.

### 2.5 Split-sample and out-of-sample validation of multimodal fusion

For validation of the multimodal fusion approach, a split-sample validation analysis was performed with similar age distributions. This was achieved by splitting the original sample based on odd/even parity of ranked participants’ ages. Linked ICA using a dimensionality of 40 with 1000 iterations (same as the CC280 main analysis) was performed on each sub-sample. Linked ICA components of the main sample were spatially correlated with linked ICA components of the validation sub-samples to examine the robustness of linked ICA to characterize brain patterns in independent components. Each of linked ICA components in one sample was matched to the component that showed the highest correlation with it in the comparing sample.

To further assess the reliability of fusion between neuroimaging modalities using the linked ICA approach in a larger sample size, linked ICA was performed in an independent sample using the same processing steps and settings. This sample was the CC420 as described above in 2.1, a subset of the Cam-CAN cohort comprised of participants who were included in CC700 but were not included in CC280 main analysis because they were either not selected to enter CC280 or had missing data from CC280. CC420 lacked ASL data so the linked ICA included 6 inputs only (DMN, SN, right FPN, left FPN, RSFA, and GMV). Other protocols were the same as the main analysis (i.e., the acquisition and preprocessing of neuroimaging data, functional network decomposition using group-ICA, and multimodal fusion using linked ICA).

### 2.6 Statistical analysis

Demographic variables were compared between age groups using one-way ANOVA for continuous variables and using chi-square test for categorical variable. Matching between functional network spatial maps and corresponding network templates was analyzed using spatial correlation tests.

To investigate the relationship between linked ICA subject loadings of each component with cognition and how it varied with age, component subject loadings from linked ICA output were analyzed in relation to Cattell test total score using regression analysis with robust fitting algorithm (Matlab function fitlm.m). Each linked ICA component subject loading score was used as the dependent variable. Cattell score, age, their interaction (age*Cattell) and quadratic effect of age were used as the independent variables in the model. Covariates of no interest included gender and head motion. The model’s formula took the following form using Wilkinson notation (Wilkinson & Rogers, 1973): IC ∼ Cattell*age + age^2 + gender + head motion. To investigate whether variance in head size across subjects could explain biologically plausible effects in addition to confounding effects, we performed an additional regression analysis with total intracranial volume (TIV) as a covariate along with the other covariates mentioned above (IC ∼ Cattell*age + age^2 + gender + head motion + TIV). The overall model fit was corrected for multiple comparison using the Bonferroni correction of family-wise error rate (FWER). A corrected *P* < 0.05 was chosen as the significance level. Only those models with significant overall model fit after FWER-correction were considered as relevant in this study. Components that were not significant after FWER-correction were considered as components not related to the predictors in the models, but possibly related to other factors such as noise signals. All statistical analyses were performed in Matlab version 2020b.

## 3. RESULTS

### 3.1 Characteristics of participants

The demographic characteristics of participants are reported in Table 1. Performance on the Cattell test was highly correlated with age within the CC280 main sample (R = -0.64, *P* < 0.0001) and CC420 validation sample (R = -0.69, *P* < 0.0001) (Figure 2).

**Figure 2.**
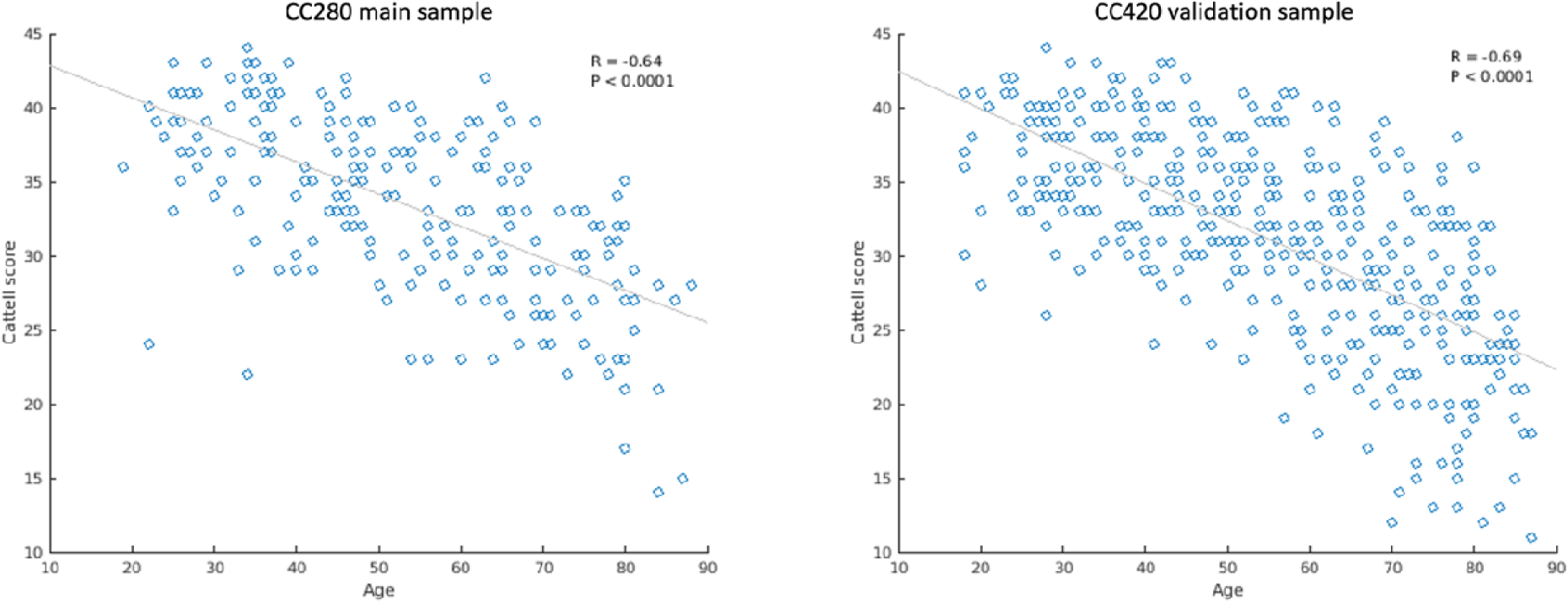
Scatter plots showing the correlation between age and fluid intelligence measured by Cattell test score in the CC280 main sample (*n* = 215) and CC420 validation sample (*n* = 433).

**Table 1.** Characteristics of participants.

| | Age range | | | | | | | | Difference<br>between<br>deciles<br>(ANOVA or $\chi^2$) |
| --- | --- | --- | --- | --- | --- | --- | --- | --- | --- |
|  | All | 18-27 | 28-37 | 38-47 | 48-57 | 58-67 | 68-77 | 78-88 | P-values |
| **CC280** |  |  |  |  |  |  |  |  |  |
| *n* | 215 | 17 | 38 | 36 | 35 | 38 | 26 | 25 |  |
| Mean age (years) | 52.9 | 24.6 | 33.5 | 43.6 | 52.2 | 62.8 | 72.3 | 81.0 |  |
| Gender |  |  |  |  |  |  |  |  |  |
| *n* (%) |  |  |  |  |  |  |  |  |  |
| Males | 106 (49.3) | 7 (41.2) | 19 (50) | 19 (52.8) | 18 (51.4) | 19 (50) | 12 (46.2) | 12 (48) | 0.99 |
| Females | 109 (50.7) | 10 (58.8) | 19 (50) | 17 (47.2) | 17 (48.6) | 19 (50) | 14 (53.8) | 13 (52) |  |
| Cattell score |  |  |  |  |  |  |  |  |  |
| Mean ± SD | 33.5±6.0 | 37.8±4.4 | 38.4±4.5 | 35.5±3.8 | 33.6±4.5 | 32.2±5.0 | 28.9±4.2 | 26.4±5.7 | < 0.0001 |
| Mini-Mental State Examination |  |  |  |  |  |  |  |  |  |
| Mean ± SD | 29.2±1.0 | 29.3±0.9 | 29.7±0.6 | 29.1±1.2 | 29.3±0.8 | 29.1±1.0 | 29.0±1.2 | 28.7±1.4 | 0.0099 |
| **CC420** |  |  |  |  |  |  |  |  |  |
| *n* | 433 | 34 | 66 | 62 | 64 | 62 | 75 | 70 |  |
| Mean age (years) | 55.1 | 22.8 | 32.4 | 42.5 | 52.5 | 62.3 | 72.1 | 81.3 |  |
| Gender |  |  |  |  |  |  |  |  |  |
| *n* (%) |  |  |  |  |  |  |  |  |  |
| Males | 212 (49.0) | 16 (47.1) | 31 (47.0) | 28 (45.2) | 31 (48.4) | 31 (50.0) | 38 (50.7) | 37 (52.9) | 0.98 |
| Females | 221 (51.0) | 18 (52.9) | 35 (53.0) | 34 (54.8) | 33 (51.6) | 31 (50.0) | 37 (49.3) | 33 (47.1) |  |
| Cattell score |  |  |  |  |  |  |  |  |  |
| Mean ± SD | 31.0±7.0 | 37.3±3.7 | 36.5±4.0 | 34.9±4.5 | 33.4±4.6 | 29.6±5.3 | 26.4±6.2 | 23.5±5.6 | < 0.0001 |
| Mini-Mental State Examination |  |  |  |  |  |  |  |  |  |
| Mean ± SD | 28.8±1.3 | 29.1±1.5 | 29.4±1.1 | 29.1±1.1 | 29.1±1.2 | 29.0±1.2 | 28.4±1.3 | 27.9±1.5 | < 0.0001 |

### 3.2 Group-average effects on functional networks, RSFA, CBF and grey matter maps

Among the 15 components generated from group-ICA, whole brain spatial maps associated with the following networks of interest specified a priori were identified: the DMN, the SN, and the lateralized FPNs (Figure 3a). The correlation between each functional network spatial map and its corresponding template from a previous study (Shirer et al., 2012) was *r* = 0.62, *P* < 0.0001 for the DMN, *r* = 0.58, *P* < 0.0001 for the SN, *r* = 0.55, *P* < 0.0001 for the right FPN, and *r* = 0.54, *P* < 0.0001 for the left FPN.

**Figure 3.**
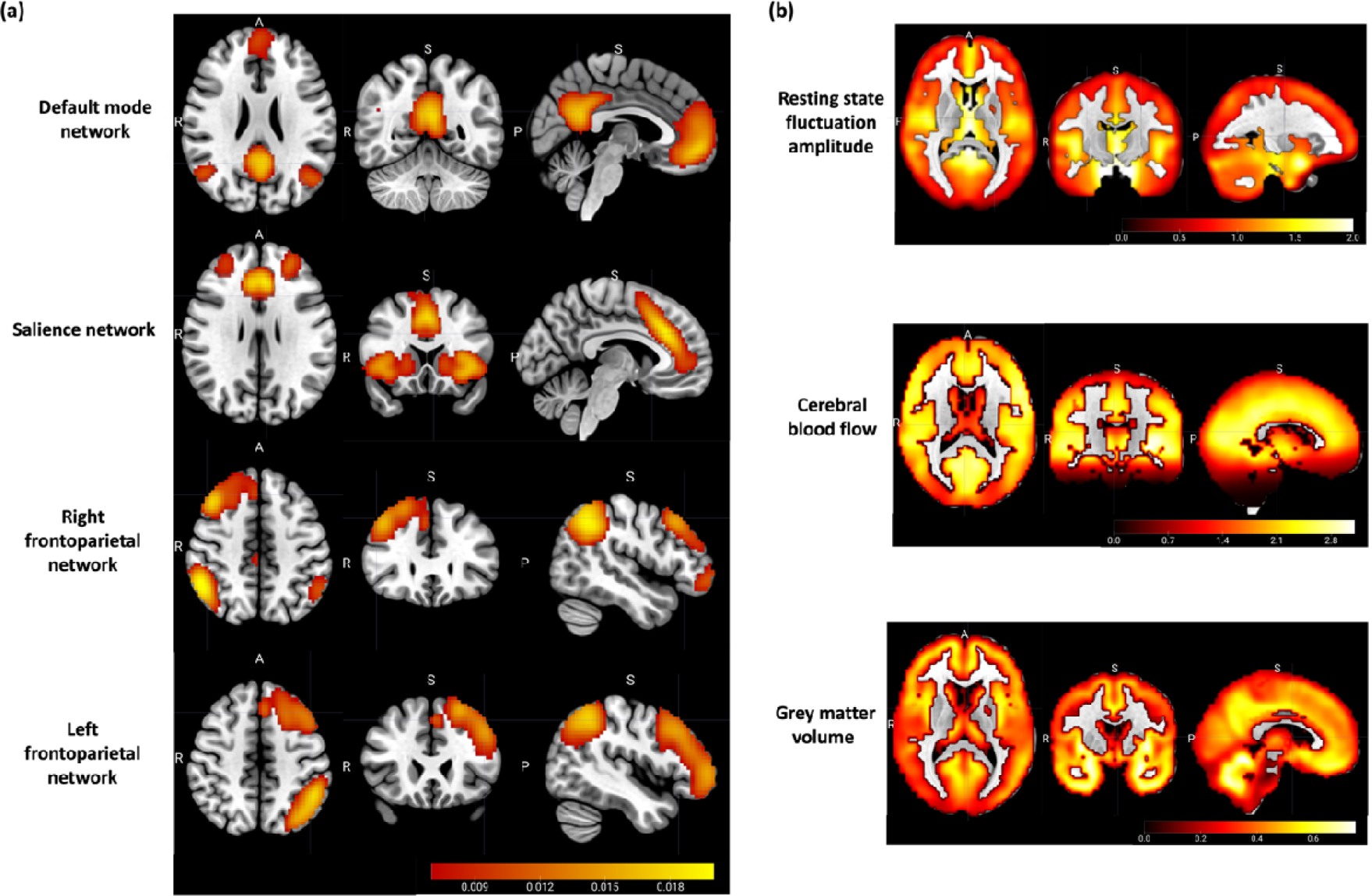
(a) The group-average spatial maps associated with the default mode network, the salience network, and the lateralized frontoparietal networks, generated from group-level independent component analysis of 648 subjects from Cam-CAN cohort Stage 2. (b) The group-average spatial maps of cerebrovascular and structural neuroimaging modality inputs to linked ICA, including resting state fluctuation amplitude and cerebral blood flow as cerebrovascular measurements, and grey matter volume as a structural measurement.

CC280 group-average spatial maps of RSFA, CBF, and GMV that were entered into linked ICA are shown in Figure 3b. Relatively strong group-average RSFA signals were observed in the temporal lobe and subcortical regions; relatively high CBF was observed in cortical and subcortical regions including frontal, posterior cingulate, pericalcarine, temporal, insula and thalamic regions; and relatively high GMV was observed in the temporal lobe, prefrontal, middle and superior frontal areas, medial occipital areas, and cerebellum.

### 3.3 Multimodal fusion using linked ICA

The relative weight of modalities in each component of CC280 is shown in Figure 4. Only modalities with significant weight (i.e., pseudo-Z-score > 3.34 which corresponds to *P* < 0.001) are presented. Two out of 40 components were excluded due to no values beyond the significance threshold from any modality. Most components (> 75%) were dominated by a single input neuroimaging modality. Components reflecting structural and cerebrovascular inputs explained overall more variance compared to resting-state functional network topography. Fusion in the same component between imaging inputs were observed between GMV, CBF and RSFA maps (i.e., IC1, IC4, IC14, IC33). Fusion was also observed between different functional networks (i.e., IC19, IC24, IC26, IC38). However, no significant fusion was observed between functional network, cerebrovascular and structural spatial maps.

**Figure 4.**
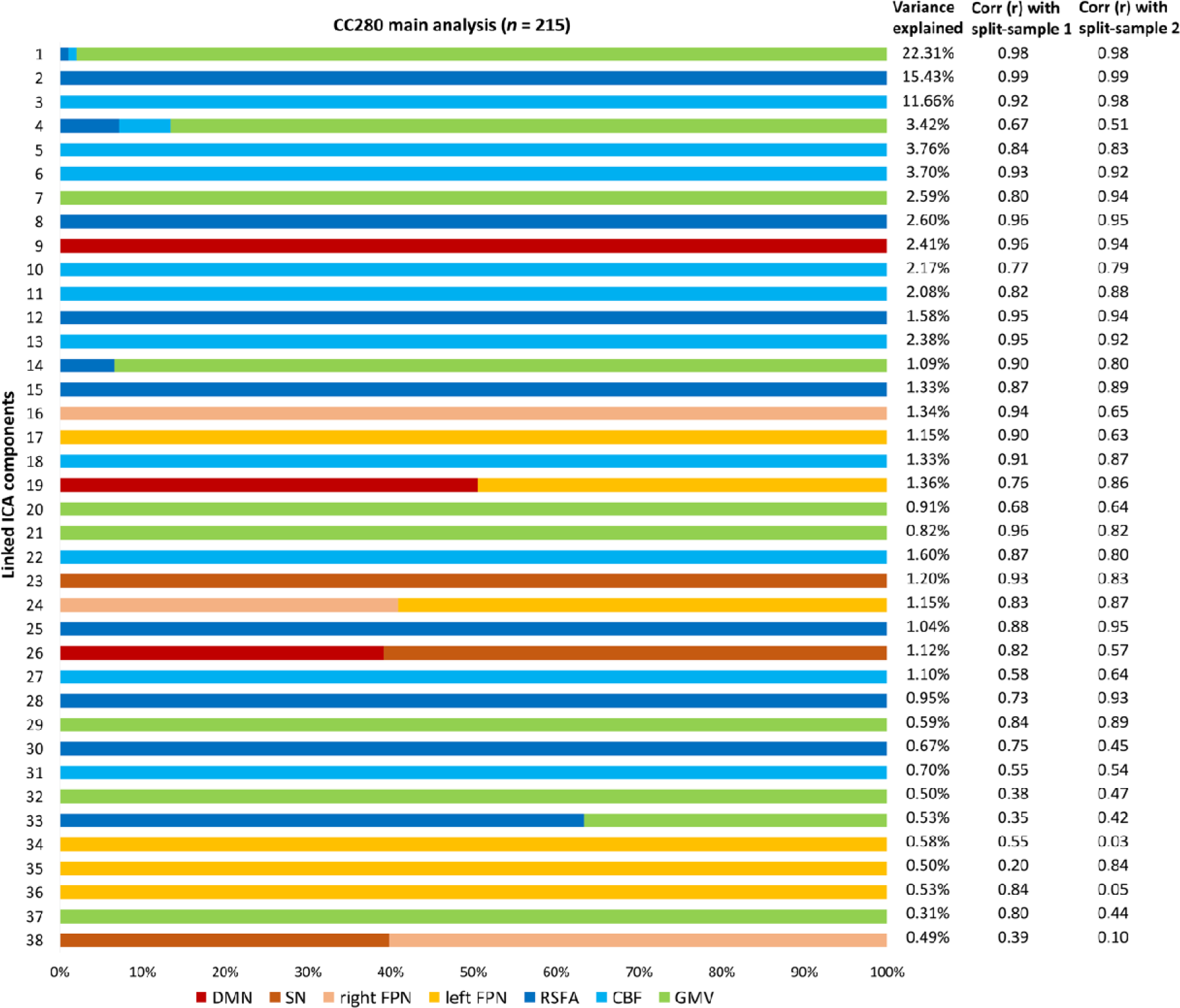
The relative weight of modalities in each component generated from linked independent component analysis (ICA) and the percentage of variance explained of each component of the CC280 main analysis (*n* = 215). Note that most components are dominated by one modality. Two columns on the right show the spatial correlation coefficients between each of linked ICA components of the CC280 main sample and split-sample validation sub-sample 1 (split-sample 1, n = 108), and main sample and split-sample validation sub-sample 2 (split-sample 2, n = 107). Abbreviations: DMN, default mode network; SN, salience network; FPN, frontoparietal network; RSFA, resting state fluctuation amplitude; CBF, cerebral blood flow; GMV, grey matter volume.

For the components considered relevant in this study (i.e., components with significant overall model fit after FWER-correction in regression), the spatial patterns of the split-sample validation analysis were generally similar to those of the main analysis, as shown by the spatial correlation between the linked ICA components of the CC280 main sample and split-sample validation sub-samples (Figure 4). The average spatial correlation of the relevant components was *r* = 0.83 between the main sample and sub-sample 1, *r* = 0.77 between the main sample and sub-sample 2, and *r* = 0.76 between sub-sample 1 and sub-sample 2.

Linked ICA was repeated with 30 and 50 numbers of components in order to ensure the results were not significantly affected by the ICA dimensionality. Fusion patterns between modalities were consistent across different model orders (Figure 5). The relative weight of modalities in each component of linked ICA with only one functional network input is shown in Supplementary Figure 3. No fusion was found between the functional network and cerebrovascular/structural patterns.

**Figure 5.**
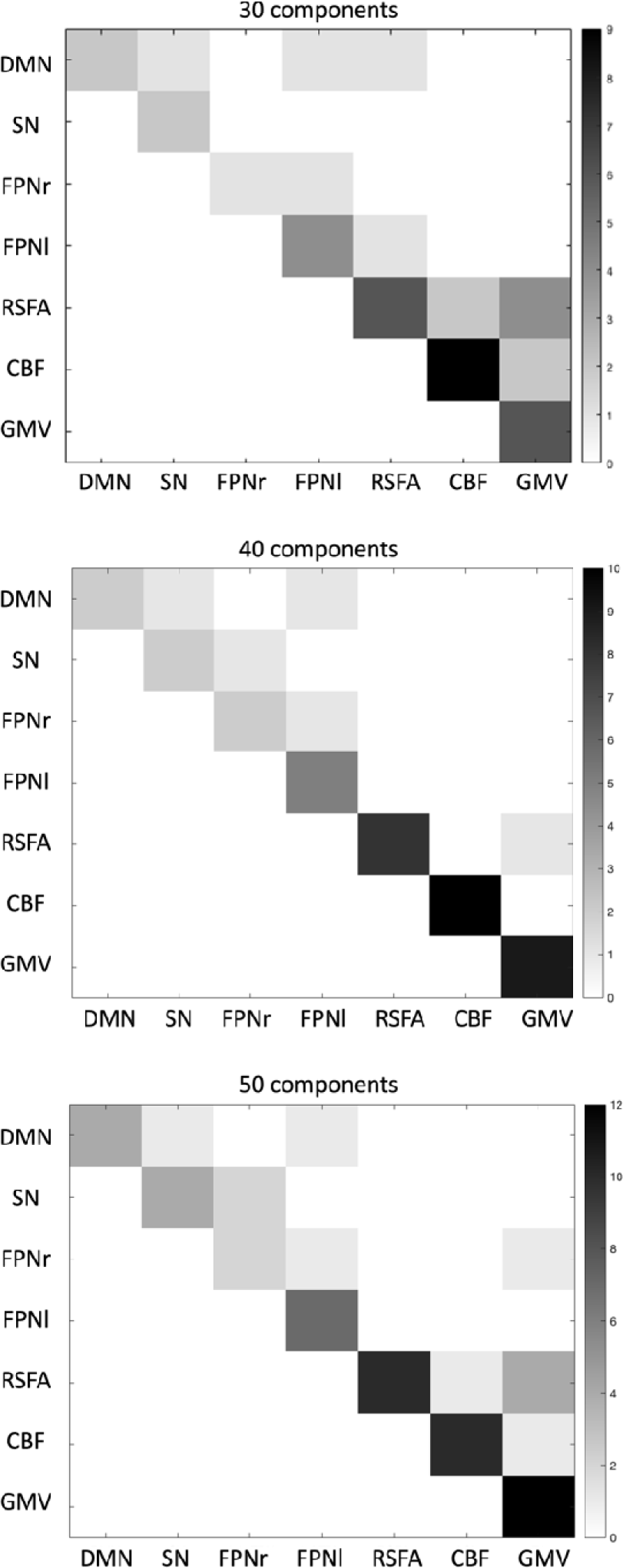
Degree of fusion between the 7 neuroimaging modalities included in linked independent component analysis (ICA) CC280 (*n* = 215) with 30, 40, and 50 components, respectively. Greater number (i.e., darker color) in the matrix represents more fusion found between the two modalities in linked ICA output components. Abbreviations: DMN, default mode network; SN, salience network; FPNr, right frontoparietal network; FPNl, left frontoparietal network; RSFA, resting state fluctuation amplitude; CBF, cerebral blood flow; GMV, grey matter volume.

### 3.4 Age- and cognition-related effects on linked ICA subject loadings

Results of regression analysis of the CC280 are shown in Table 2. The overall model fits of 15 components remained significant after FWER-correction. Components that were not significant after FWER-correction were considered as components not related to the predictors in the models, but possibly related to other factors such as noise signals. Association with age was observed in multiple components and the strongest age effects were observed in components related to GMV, CBF and RSFA (Figure 6). The strongest non-linear changes in relation to age were observed in IC4, IC7 and IC22. Among the 15 components of interest, Cattell score was positively correlated with IC1 which reflected global GMV with regional CBF and RSFA signals, IC16 which reflected the right FPN pattern and IC17 which reflected the left FPN pattern, accounting for age, gender and head motion as covariates. Spatial maps of IC1, IC16 and IC17, accompanied by scatter plots showing models of Cattell test score against IC subject loadings, are shown in Figure 7. Results of IC1 indicate that subjects with higher Cattell score had higher subject loadings indicating i) higher GMV globally; ii) higher CBF mainly in the lingual gyrus, calcarine, thalamus, and cingulate gyrus coupled with low perfusion in middle temporal gyrus; and iii) higher RSFA values in areas proximal to vascular and cerebrospinal fluid territories (venous sinuses and middle cerebral arteries).

**Figure 6.**
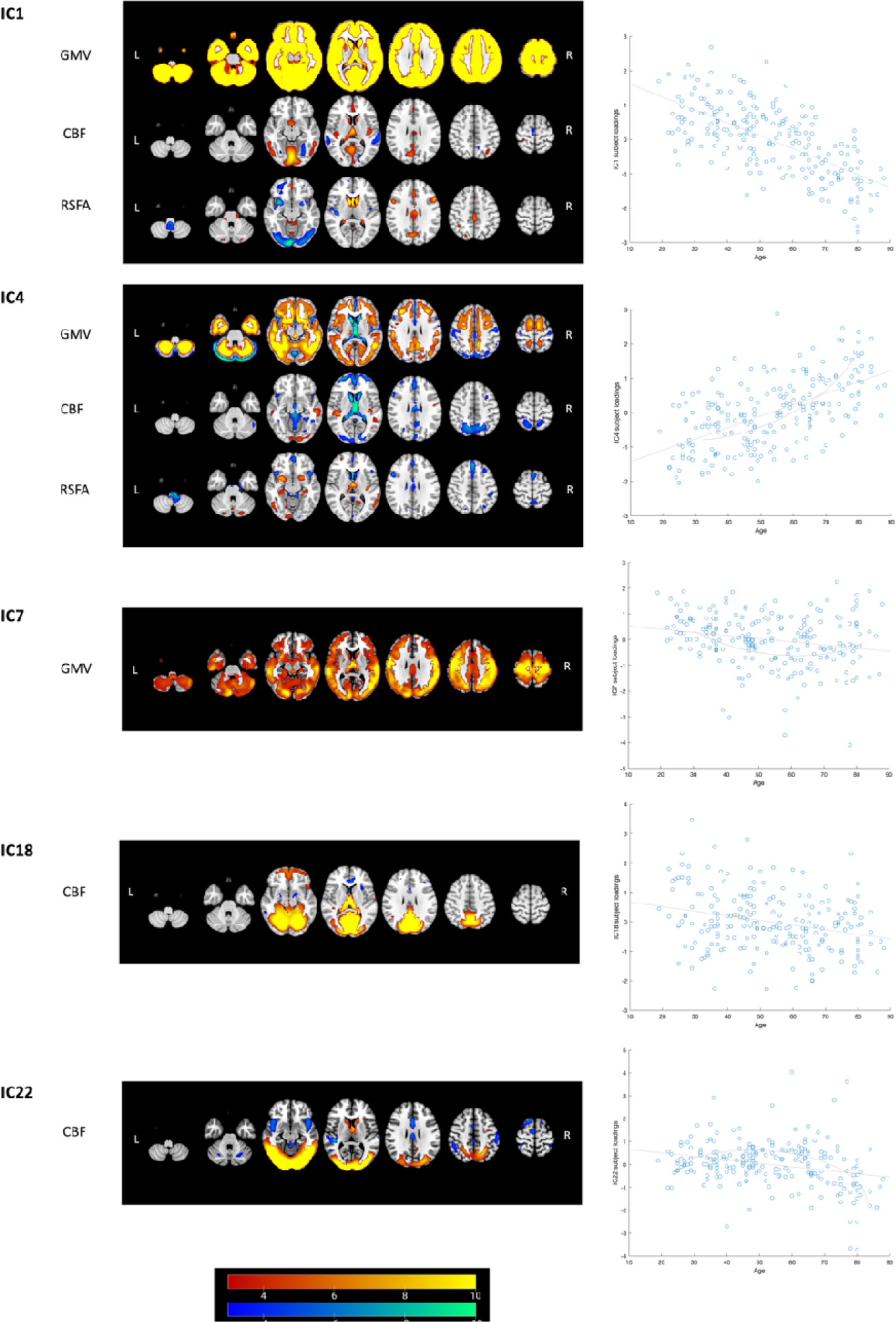
Linked ICA weighted spatial maps for five components showing strong age effects on subject loadings in CC280 main analysis (*n* = 215). Warm and cold colour scheme indicate positive and negative association with linked ICA subject loadings, respectively. For example, an individual with a high loading value on IC1, i.e., young adult, had i) high whole brain GMV, ii) high perfusion in visual cortex and posterior cingulate cortex (PCC) coupled with low perfusion in middle temporal gyrus and iii) high RSFA in dorsolateral prefrontal cortex and PCC, coupled with low RSFA values in areas proximal to vascular and cerebrospinal fluid territories (venous sinuses and middle cerebral arteries). The brain figures depict the weighted spatial maps and the accompanying scatter plots show models of age plotted against IC subject loadings. Non-linear changes in relation to age were observed in IC4, IC7 and IC22 (refer to Table 2 for regression results). For visualization the spatial map threshold is set to 3 < |Z| < 10. Abbreviations: CBF, cerebral blood flow; GMV, grey matter volume; RSFA, resting state fluctuation amplitude.

**Figure 7.**
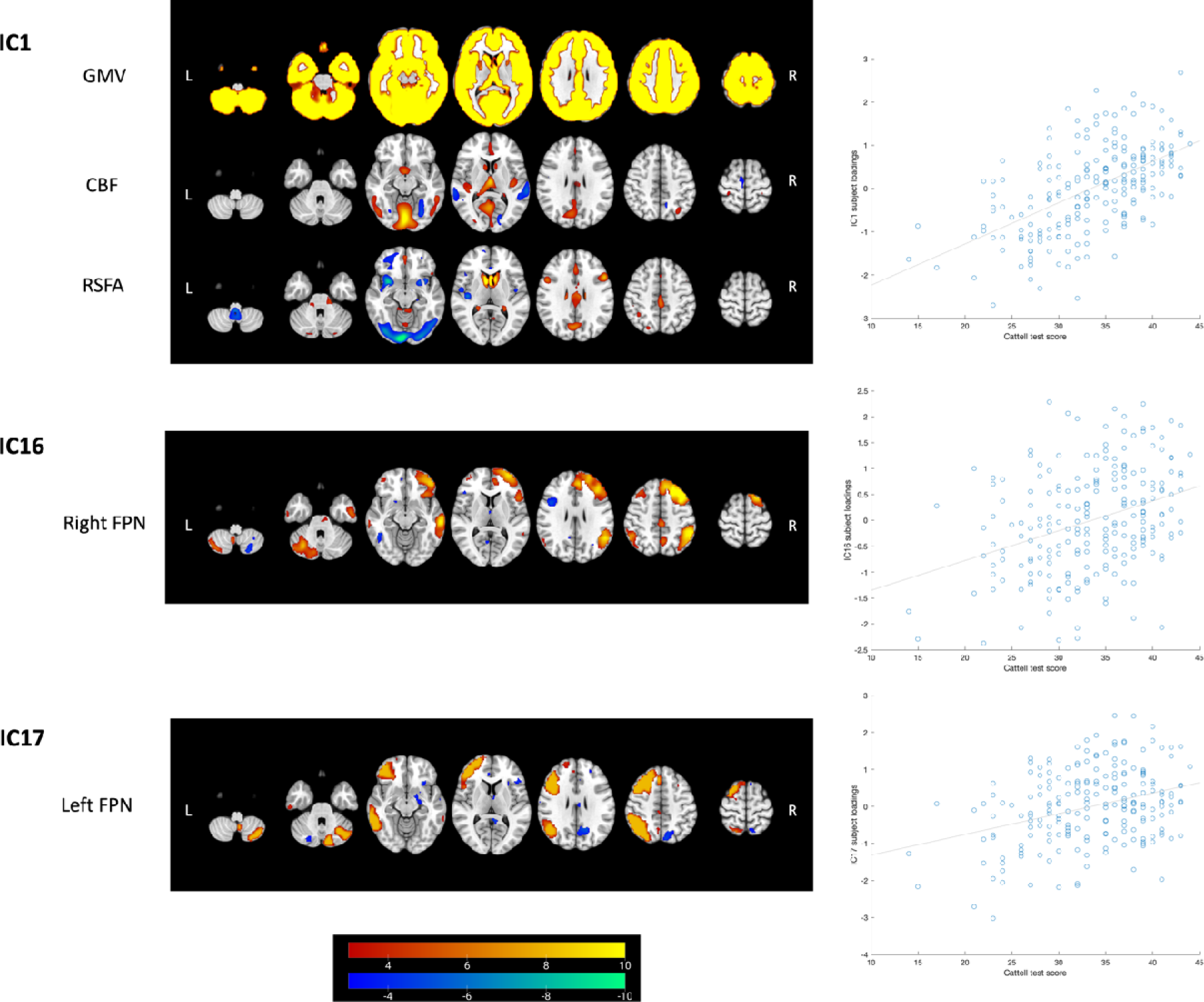
Linked ICA weighted spatial maps for three components showing unique associations with Cattell test score in CC280 main analysis (*n* = 215). Warm and cold colour scheme indicate positive and negative association with linked ICA subject loadings, respectively. The accompanying scatter plots show models of Cattell test score plotted against IC subject loadings. One component reflects signals from structural and cerebrovascular measurements: IC1 which reflects grey matter volume (GMV), cerebral blood flow (CBF) and resting state fluctuation amplitude (RSFA) (see Figure 6 and main text for further interpretation). Two components reflect signals from functional networks: IC16 which reflects the right frontoparietal network (FPN) and IC17 which reflects the left FPN. For visualization the spatial map threshold is set to 3 < |Z| < 10. Similar components of IC1 and IC16 were found in CC420 out-of-sample validation analysis (supplementary materials) to be associated with fluid intelligence.

**Table 2.**
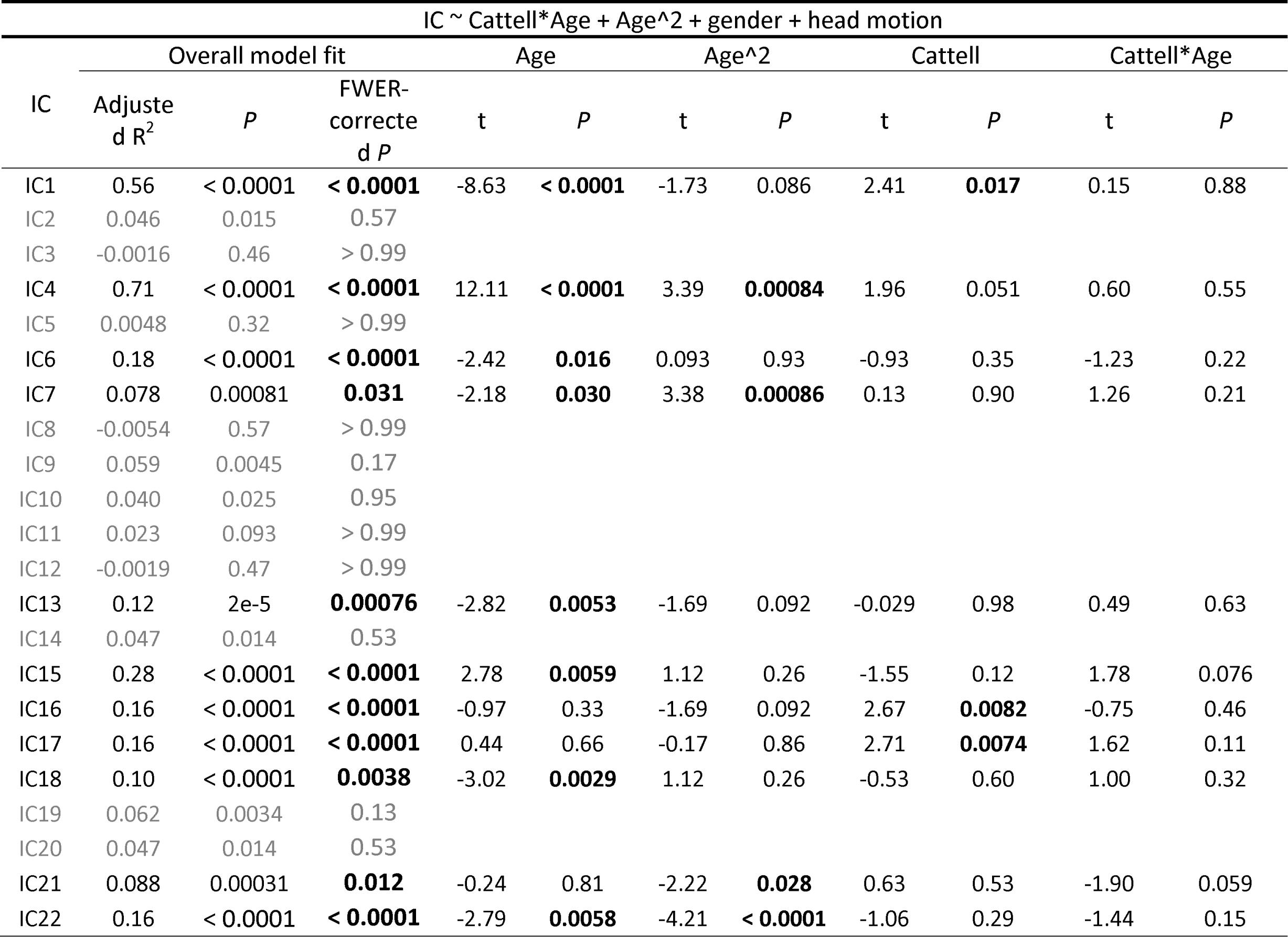
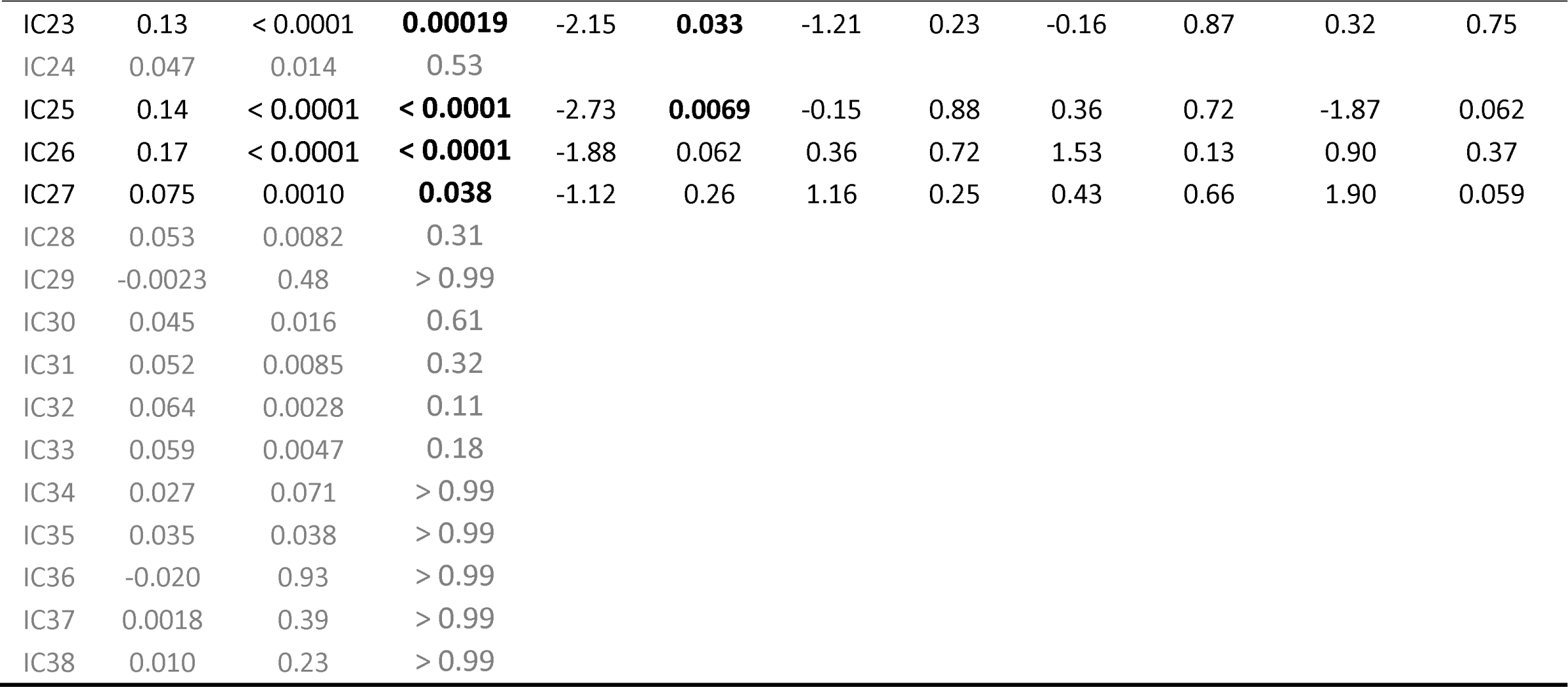
Multiple regression analysis results of each independent component (IC) subject loadings from linked independent component analysis of CC280 participants (38 components based on 7 modalities and n = 215).

| IC ~ Cattell*Age + Age^2 + gender + head motion |  |  |  |  |  |  |  |  |  |  |  |
| --- | --- | --- | --- | --- | --- | --- | --- | --- | --- | --- | --- |
| IC | Overall model fit |  |  | Age |  | Age^2 |  | Cattell |  | Cattell*Age |  |
| | Adjusted $R^2$ | $P$ | FWER-corrected $P$ | t | $P$ | t | $P$ | t | $P$ | t | $P$ |
| IC1 | 0.56 | < 0.0001 | < 0.0001 | -8.63 | < 0.0001 | -1.73 | 0.086 | 2.41 | **0.017** | 0.15 | 0.88 |
| IC2 | 0.046 | 0.015 | 0.57 |  |  |  |  |  |  |  |  |
| IC3 | -0.0016 | 0.46 | > 0.99 |  |  |  |  |  |  |  |  |
| IC4 | 0.71 | < 0.0001 | < 0.0001 | 12.11 | < 0.0001 | 3.39 | **0.00084** | 1.96 | 0.051 | 0.60 | 0.55 |
| IC5 | 0.0048 | 0.32 | > 0.99 |  |  |  |  |  |  |  |  |
| IC6 | 0.18 | < 0.0001 | < 0.0001 | -2.42 | **0.016** | 0.093 | 0.93 | -0.93 | 0.35 | -1.23 | 0.22 |
| IC7 | 0.078 | 0.00081 | **0.031** | -2.18 | **0.030** | 3.38 | **0.00086** | 0.13 | 0.90 | 1.26 | 0.21 |
| IC8 | -0.0054 | 0.57 | > 0.99 |  |  |  |  |  |  |  |  |
| IC9 | 0.059 | 0.0045 | 0.17 |  |  |  |  |  |  |  |  |
| IC10 | 0.040 | 0.025 | 0.95 |  |  |  |  |  |  |  |  |
| IC11 | 0.023 | 0.093 | > 0.99 |  |  |  |  |  |  |  |  |
| IC12 | -0.0019 | 0.47 | > 0.99 |  |  |  |  |  |  |  |  |
| IC13 | 0.12 | 2e-5 | **0.00076** | -2.82 | **0.0053** | -1.69 | 0.092 | -0.029 | 0.98 | 0.49 | 0.63 |
| IC14 | 0.047 | 0.014 | 0.53 |  |  |  |  |  |  |  |  |
| IC15 | 0.28 | < 0.0001 | < 0.0001 | 2.78 | **0.0059** | 1.12 | 0.26 | -1.55 | 0.12 | 1.78 | 0.076 |
| IC16 | 0.16 | < 0.0001 | < 0.0001 | -0.97 | 0.33 | -1.69 | 0.092 | 2.67 | **0.0082** | -0.75 | 0.46 |
| IC17 | 0.16 | < 0.0001 | < 0.0001 | 0.44 | 0.66 | -0.17 | 0.86 | 2.71 | **0.0074** | 1.62 | 0.11 |
| IC18 | 0.10 | < 0.0001 | **0.0038** | -3.02 | **0.0029** | 1.12 | 0.26 | -0.53 | 0.60 | 1.00 | 0.32 |
| IC19 | 0.062 | 0.0034 | 0.13 |  |  |  |  |  |  |  |  |
| IC20 | 0.047 | 0.014 | 0.53 |  |  |  |  |  |  |  |  |
| IC21 | 0.088 | 0.00031 | **0.012** | -0.24 | 0.81 | -2.22 | **0.028** | 0.63 | 0.53 | -1.90 | 0.059 |
| IC22 | 0.16 | < 0.0001 | < 0.0001 | -2.79 | **0.0058** | -4.21 | < 0.0001 | -1.06 | 0.29 | -1.44 | 0.15 |
| IC23 | 0.13 | < 0.0001 | **0.00019** | -2.15 | **0.033** | -1.21 | 0.23 | -0.16 | 0.87 | 0.32 | 0.75 |
| IC24 | 0.047 | 0.014 | 0.53 |  |  |  |  |  |  |  |  |
| IC25 | 0.14 | < 0.0001 | **< 0.0001** | -2.73 | **0.0069** | -0.15 | 0.88 | 0.36 | 0.72 | -1.87 | 0.062 |
| IC26 | 0.17 | < 0.0001 | **< 0.0001** | -1.88 | 0.062 | 0.36 | 0.72 | 1.53 | 0.13 | 0.90 | 0.37 |
| IC27 | 0.075 | 0.0010 | **0.038** | -1.12 | 0.26 | 1.16 | 0.25 | 0.43 | 0.66 | 1.90 | 0.059 |
| IC28 | 0.053 | 0.0082 | 0.31 |  |  |  |  |  |  |  |  |
| IC29 | -0.0023 | 0.48 | > 0.99 |  |  |  |  |  |  |  |  |
| IC30 | 0.045 | 0.016 | 0.61 |  |  |  |  |  |  |  |  |
| IC31 | 0.052 | 0.0085 | 0.32 |  |  |  |  |  |  |  |  |
| IC32 | 0.064 | 0.0028 | 0.11 |  |  |  |  |  |  |  |  |
| IC33 | 0.059 | 0.0047 | 0.18 |  |  |  |  |  |  |  |  |
| IC34 | 0.027 | 0.071 | > 0.99 |  |  |  |  |  |  |  |  |
| IC35 | 0.035 | 0.038 | > 0.99 |  |  |  |  |  |  |  |  |
| IC36 | -0.020 | 0.93 | > 0.99 |  |  |  |  |  |  |  |  |
| IC37 | 0.0018 | 0.39 | > 0.99 |  |  |  |  |  |  |  |  |
| IC38 | 0.010 | 0.23 | > 0.99 |  |  |  |  |  |  |  |  |

Among the components of interest, TIV altered the correlation with Cattell on IC1 (i.e., the Cattell coefficient was no longer significant, *t* = -0.19, *P* = 0.85, by including TIV as a covariate in the regression model). This suggests that individuals with high TIV and high Cattell performance expressed strongly IC1. No evidence was found for TIV explaining the effects of age on subject loadings across all ICs.

### 3.5 Multimodal fusion using linked ICA in an independent Cam-CAN subset

Results of the out-of-sample validation analysis using CC420 were reported in supplementary materials. Major results were consistent with the main analysis.

## 4. DISCUSSION

We present a multivariate data-driven analysis of the patterns of structural, cerebrovascular and functional change in the brain across the adult lifespan in healthy subjects from 18 – 88 years old. The main results are that (i) there were concordant changes in morphometry and cerebrovascular signals, but not between resting-state functional network spatial maps and morphometry or cerebrovascular signals; and (ii) the variance in expression of linked ICA components was cognitively relevant after adjusting for age and other covariates of no interest. In particular, individual differences in fluid intelligence correlated with (i) diffuse brain atrophy coupled with regional cerebrovascular differences and (ii) resting-state network activity in the right FPN. These principal findings were replicated in an independent cohort, without ASL data, and in split-sample analysis of the original cohort with ASL data. We present the insights from linked ICA to bring together measurements from multimodal neuroimaging with their independent and additive information to characterize structural, functional and cerebrovascular brain changes of healthy ageing.

The results demonstrate the robustness of linked ICA to characterize brain patterns comprising information from multiple neuroimaging measurements in independent components by repeating linked ICA in different sample sizes and dimensionalities. The independently distributed structural-cerebrovascular and functional patterns underline the need for a precise method to integrate information from multiple neuroimaging measurements in order to comprehensively characterize brain pattern and associated variability in cognition.

The linked ICA identified a strong structural effect, in that the component showing global GMV change (i.e., IC1 from the main analysis) explained the most variance. This is consistent with previous studies of ageing using linked ICA (Doan, Engvig, Zaske, et al., 2017; Douaud et al., 2014). Cerebrovascular measures were identified in the same component, suggesting that the atrophy effects were partly linked to cerebrovascular health. This accords with large-scale lifespan studies showing global brain atrophy association with cerebrovascular changes (Asllani et al., 2009; Iadecola, 2017; Kennedy & Raz, 2015; Lemaitre et al., 2012; Peelle, Cusack, & Henson, 2012). Strongest changes with age were also observed in components reflecting GMV and CBF. Most age effects were linear while four components showed statistically significant quadratic changes, consistent with current literature showing significant age-related alternations in GMV and cerebrovascular activity (Bethlehem et al., 2022; Kievit et al., 2014; K. A. Tsvetanov, Henson, Jones, et al., 2021). In contrast with atrophy and cerebrovascular indices, there was little fusion in the output components between resting-state functional networks and other modalities. A previous study including the DMN in linked ICA together with grey matter density, area, thickness, mean diffusivity, fractional anisotropy, and radial diffusivity also showed little fusion between the DMN and other modalities in the output components (Maglanoc et al., 2020). The distributed structural, cerebrovascular and functional topography warrants integrated and multimodal neuroimaging analytical approach, as these neuroimaging measurements could indicate independent and additive information about ageing and cognition.

Fluid intelligence is the core of psychometric analyses of intelligence and correlated with other cognitive tests including tests that assess successful day-to-day functioning in society (Ghisletta et al., 2012; Marsiske & Willis, 1995). Within our sample, we assessed fluid intelligence using the Cattell task. Performance on this task declined with age, consistent with previously demonstrated negative correlation with age in both cross-sectional (Hartshorne & Germine, 2015; Kievit et al., 2016; Kievit et al., 2014) and longitudinal studies (Ghisletta et al., 2012; Timothy A. Salthouse, 2010; Schaie, 1994). Multiple regression analysis using the linked ICA component subject loadings indicates that global GMV coupled with regional cerebrovascular activity (IC1) and the right FPN (IC16) were positively correlated with fluid intelligence after adjusting for age and other covariates of no interest. Results of IC1 indicate that subjects with higher fluid intelligence had i) higher GMV globally; ii) higher CBF mainly in the lingual gyrus, calcarine, thalamus, and cingulate gyrus coupled with low perfusion in middle temporal gyrus; and iii) higher RSFA values in areas proximal to vascular and cerebrospinal fluid territories. The paradoxical hypoperfusion in middle frontal gyrus for young adults and high performers may be explained by higher perfusion values in old adults or poor performers. Macrovascular artifacts are common in ASL findings (Detre, Rao, Wang, Chen, & Wang, 2012; H. J. Mutsaerts et al., 2017; K. A. Tsvetanov, Henson, Jones, et al., 2021) due to prolonged arterial transfer times with ageing (Dai et al., 2017). The increase of RSFA with age or poor cognition in vascular regions is consistent with previous studies (Makedonov et al., 2013; Theyers, Goldstein, Metcalfe, Robertson, & MacIntosh, 2019; K. A. Tsvetanov et al., 2015; K. A. Tsvetanov, Henson, Jones, et al., 2021; Kamen A Tsvetanov et al., 2022; Viessmann, Moller, & Jezzard, 2019), which likely reflects pulsatile signals known to increase with atherosclerosis and vessel stiffening in ageing (Webb et al., 2012). The patterns reflected by IC1 and IC16 were consistently found in an independent and larger sample and had significant correlations with fluid intelligence. TIV accounted for most of the correlation between IC1 and fluid intelligence, consistent with previous findings on the link between head size and fluid intelligence (Lee, McGue, Iacono, Michael, & Chabris, 2019). The FPN is an important control network, in which functional integration is positively correlated with general cognitive ability including fluid intelligence (Marek & Dosenbach, 2018; Samu et al., 2017; Sheffield et al., 2015). The current results are compatible with previous reports that the across-network connectivity of resting-state FPN is positively correlated with fluid intelligence (Bethlehem et al., 2020; Cole, Ito, & Braver, 2015; Hearne, Mattingley, & Cocchi, 2016). The results also suggest that when cognitively healthy, the right FPN activity is positively correlated with fluid intelligence regardless of age.

A major advantage of linked ICA is its ability to combine imaging modalities with different spatial dimensions or features by applying ICA on each modality while accounting for the spatial correlation of each modality, enabling us to model shared variance across different imaging modalities (Groves et al., 2011; Groves et al., 2012). Hence, the derived components may be more sensitive to an effect of interest especially when the effect is present across different imaging modalities (Francx et al., 2016). Linked ICA has revealed morphological patterns that are related to age, cognition, and Alzheimer’s disease (Alnaes et al., 2018; Doan, Engvig, Persson, et al., 2017; Doan, Engvig, Zaske, et al., 2017; Douaud et al., 2014; Groves et al., 2012) and predicted brain morphological patterns in neuropsychiatric disorders such as depression (Maglanoc et al., 2020), schizophrenia (Brandt et al., 2015; Doan, Kaufmann, et al., 2017), bipolar disorders (Doan, Kaufmann, et al., 2017), and attention-deficit/hyperactivity disorder (ADHD) (Francx et al., 2016). However, many previous studies using linked ICA focused on co-modelling brain structural effects across modalities, for example combining only grey matter morphological measures (e.g., grey matter volume/density, cortical thickness) or combining grey with white matter properties (Doan, Engvig, Zaske, et al., 2017; Doan, Kaufmann, et al., 2017; Douaud et al., 2014; Francx et al., 2016). In the present study, we showed the potential to characterize joint changes in functional, cerebrovascular and structural measures and disentangle their relationships with cognition and ageing. We found no cognitively relevant fusion between functional network spatial maps and structural or cerebrovascular spatial maps. The majority of components were dominated by a single neuroimaging measurement. It suggests that variability of brain patterns in healthy ageing subjects is better characterized by multiple independent components dominated by one of the structural, cerebrovascular or functional network measurements, but not captured in a single component reflecting all of these signals. This is contrary to our hypothesis that concordant changes on functional, structural and cerebrovascular activities would be observed, as it is a common view that age-related changes in vasculature, brain anatomy and brain function are a complex interplay that affects cognition (Fabiani, Rypma, & Gratton, 2021; Zimmerman et al., 2021). Previous studies have also reported a correlation between brain functional and structural connectivity in healthy subjects (Horn, Ostwald, Reisert, & Blankenburg, 2014; Vazquez-Rodriguez et al., 2019). Nevertheless, the current results of linked ICA do not necessarily indicate no correlation between functional network activity and structural or cerebrovascular changes, but rather suggest that age-related individual variance in the brain of cognitively-healthy subjects is better characterized by independent components representing either functional network activity alone or anatomical and cerebrovascular activity.

There are other multivariate approaches that might be more robust and sensitive in discovering covariance, such as CCA and PLS (Murley et al., 2020; Murley et al., 2022; Passamonti et al., 2019; Tibon et al., 2021; K. A. Tsvetanov, Gazzina, et al., 2021; K. A. Tsvetanov et al., 2016) as well as combinations in approaches (e.g., mCCA+jICA) (V. D. Calhoun & Sui, 2016; Sui, Adali, Yu, Chen, & Calhoun, 2012). However, the linked ICA approach offers a number of advantages. First, using CCA or PLS, where subjects and voxels are entered as samples and variables (e.g., 215x90000), would result in a multi-fold increase of variables compared to samples, which may undermine stability (e.g., a rule of thumb for CCA is to have a samples:variables ratio > 5) and make the analysis susceptible to overfitting. Alternative strategies would be to introduce an additional data reduction step (e.g., principal component analysis), regularisation or pre-whitening, or transposition of the matrices. The latter increases the computational cost and constrains the spatial correspondence. Second, linked ICA does not impose constraints on the spatial overlap between modalities. Beyond the advantage of accommodating differential spatial smoothness, linked ICA also enables detection of spatially adjacent but non-overlapping signals between structural, cerebrovascular and functional modalities (e.g., atrophy or hypoperfusion in one region may lead to changes in connectivity on a network level). Third, linked ICA can identify patterns that are multi-modal or that are sparse in modalities. Many variations of CCA or PLS exist which have been mainly optimised for two datasets, while in this study we have considered 6-7 datasets. Nonetheless, using another multivariate approach to analyze the association between two imaging modalities that are subsequently linked to performance on multiple cognitive measures could be a future work to confirm the associations of specific neuroimaging measurements with cognition.

There are limitations in the present study. First, there is no standard dimensionality to be used in ICA. However, the number of components used in group-ICA and linked ICA in the present study was based on the stable and favorable dimensionality indicated by previous literature (Beckmann et al., 2005; Damoiseaux et al., 2006; Doan, Engvig, Zaske, et al., 2017; Doan, Kaufmann, et al., 2017; Francx et al., 2016; Groves et al., 2012). Moreover, linked ICA was repeated with several dimensionalities and the results were similar: the fusion patterns in the derived components were similar and the cognitively relevant components were consistent across analyses with 30, 40 and 50 dimensions. Second, the Cattell test informed components relevant to domain-general abilities. Future research should investigate more detailed or domain-specific brain-cognition relationships. Using a variety of cognitive tests taxing different cognitive abilities enables to dissociate domain-general from domain-specific associations and better understand cognitive diversity in ageing (Shafto et al., 2020). Third, the functional network spatial maps used in linked ICA were based on associations of components with the topography of functional networks. As joint consideration of activity and connectivity might better characterize the brain dynamics and cognitive performance in normal ageing (K. A. Tsvetanov et al., 2018), it is possible that connectivity between functional nodes could indicate more information than the functional network topography alone. Future research could consider investigating the intercorrelations between functional connectivity and multiple neuroimaging modalities or integrating time-course rs-fMRI data (4D data) with 3D spatial maps from other modalities (Qi et al., 2022). Fourth, considering that we investigated healthy subjects across the whole lifespan, the relatively small sample size of the main analysis could be a limitation (Marek et al., 2022).

Advances in neuroimaging provide more insights into brain morphology, functional networks and vascular dynamics. While integration of information from multiple modalities provides more accurate representation of brain patterns, currently there are limited analysis approaches to co-model multiple neuroimaging inputs. In the present study, using linked ICA we have shown cognitively-relevant integration between grey matter and cerebrovascular changes, but minimal integration between functional networks and other modalities. The current sample comprises of cognitively healthy and, in comparison with the general population, relatively well-educated subjects. Hence, one possibility might be that for people with well-maintained cognition, resting-state functional network and structural-cerebrovascular dynamics independently characterize brain patterns that are related to age and fluid intelligence; while these would not necessarily be consistently found in cognitively impaired subjects, such as those with dementia. In subjects with dementia, the cognitively relevant dynamics of functional network, morphometry and vasculature might be more dependent on one another and such dependency might correlate to a compensation mechanism (Cabeza et al., 2018). The present study highlights the importance of future study to combine neuroimaging modalities measuring these major dynamics to characterize brain patterns related to the diagnosis and prognosis of neurodegenerative diseases.

## 5. CONCLUSION

Linked ICA can be used to integrate multiple neuroimaging modalities. We have demonstrated its ability to characterize brain pattern variability and to differentiate brain changes in healthy ageing. Across the lifespan, the most significant predictors of differences in fluid intelligence were global GMV coupled with regional cerebrovascular activity, and right FPN activity. The independently distributed structural-cerebrovascular and functional patterns in normal ageing adults underline the need for considering information from multiple neuroimaging measurements to characterize and understand brain pattern variability and cognition. Linked ICA as a multimodal neuroimaging analysis method can provide new insights into the relative brain structural, functional and vascular contributors to cognitive impairment in disorders associated with ageing, including dementia and other neurodegenerative disease.

## Data availability statement

Data of the Cambridge Centre for Ageing and Neuroscience (Cam-CAN) study are available via the Cam-CAN’s portal (https://camcan-archive.mrc-cbu.cam.ac.uk/dataaccess). Data and codes that support findings of this study are available upon reasonable request. Restrictions may apply in order to preserve participant confidentiality.

## Acknowledgments

The Cam-CAN research was supported by the Biotechnology and Biological Sciences Research Council (grant number BB/H008217/1).This work is supported by the Guarantors of Brain (G101149), the Wellcome Trust (103838), the Medical Research Council (SUAG/092 G116768; and SUAG/010 RG91365), European Union’s Horizon 2020 (732592) and the NIHR Cambridge Biomedical Research Centre (BRC-1215-20014). X.L. is supported by the Cambridge Commonwealth, European and International Trust and the China Scholarship Council. For the purpose of open access, the author has applied a CC BY public copyright licence to any Author Accepted Manuscript version arising from this submission. The views expressed are those of the authors and not necessarily those of the NHS, the NIHR or the Department of Health and Social Care.

## Ethics approval statement

Ethical approval of the Cam-CAN study was obtained from the Cambridge 2 Research Ethics Committee.

## Patient consent statement

Written informed consent was given by all participants.

