## supplementary materials for "Multimodal fusion analysis of functional, cerebrovascular and structural neuroimaging in healthy ageing subjects"

**3.5 Multimodal fusion using linked ICA in an independent Cam-CAN subset**

The relative weight of modalities in each linked ICA output component of CC420 is shown in **Supplementary** **Figure 1**. Only modalities with significant weight (i.e., pseudo-Z-score > 3.34) are presented. The majority of components (60%) were dominated by a single input neuroimaging modality.

Results of regression analysis are shown in **Supplementary** **Table**. The overall model fits of 21 components remained significant after FWER-correction and therefore these components were considered as relevant in this study. Among the 21 components of interest, including age, gender, and head motion as covariates, Cattell score was positively correlated with IC2 which reflected global GMV with regional RSFA signals and this component was similar to IC1 in CC280 analysis. An interaction between Cattell score and age was found in IC7 which reflected right FPN signals, and this component was similar to IC16 in CC280 analysis. Spatial maps of IC2 and IC7, accompanied by scatter plots showing models of Cattell test score against IC subject loadings, are shown in **Supplementary** **Figure 2**.

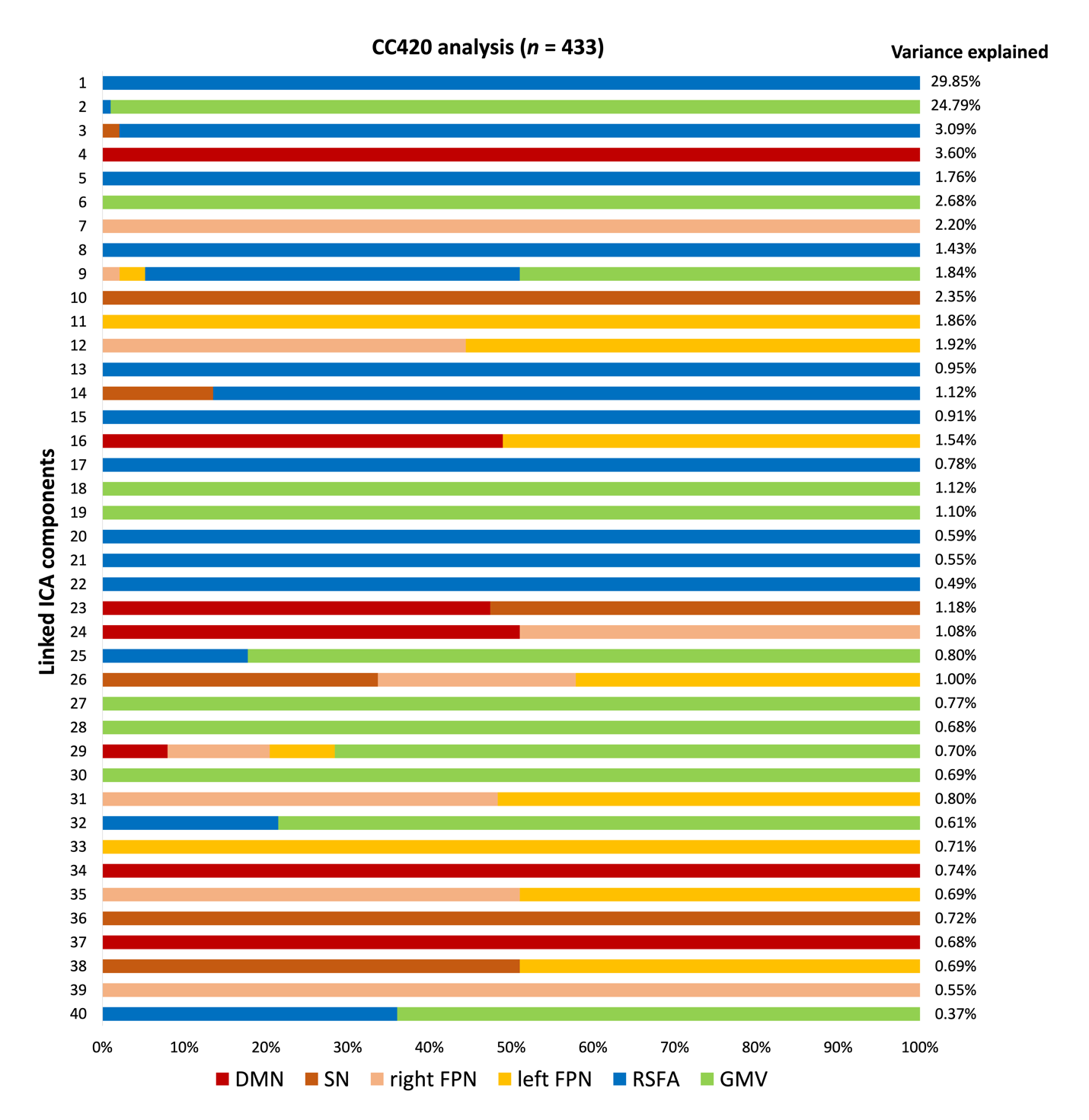

**Supplementary Figure 1.** The relative weight of modalities in each component generated from linked independent component analysis (ICA) and the percentage of variance explained of each component of the CC420 out-of-sample validation analysis (*n* = 433). Note that most components are dominated by one modality. Abbreviations: DMN, default mode network; SN, salience network; FPN, frontoparietal network; RSFA, resting state fluctuation amplitude; GMV, grey matter volume.

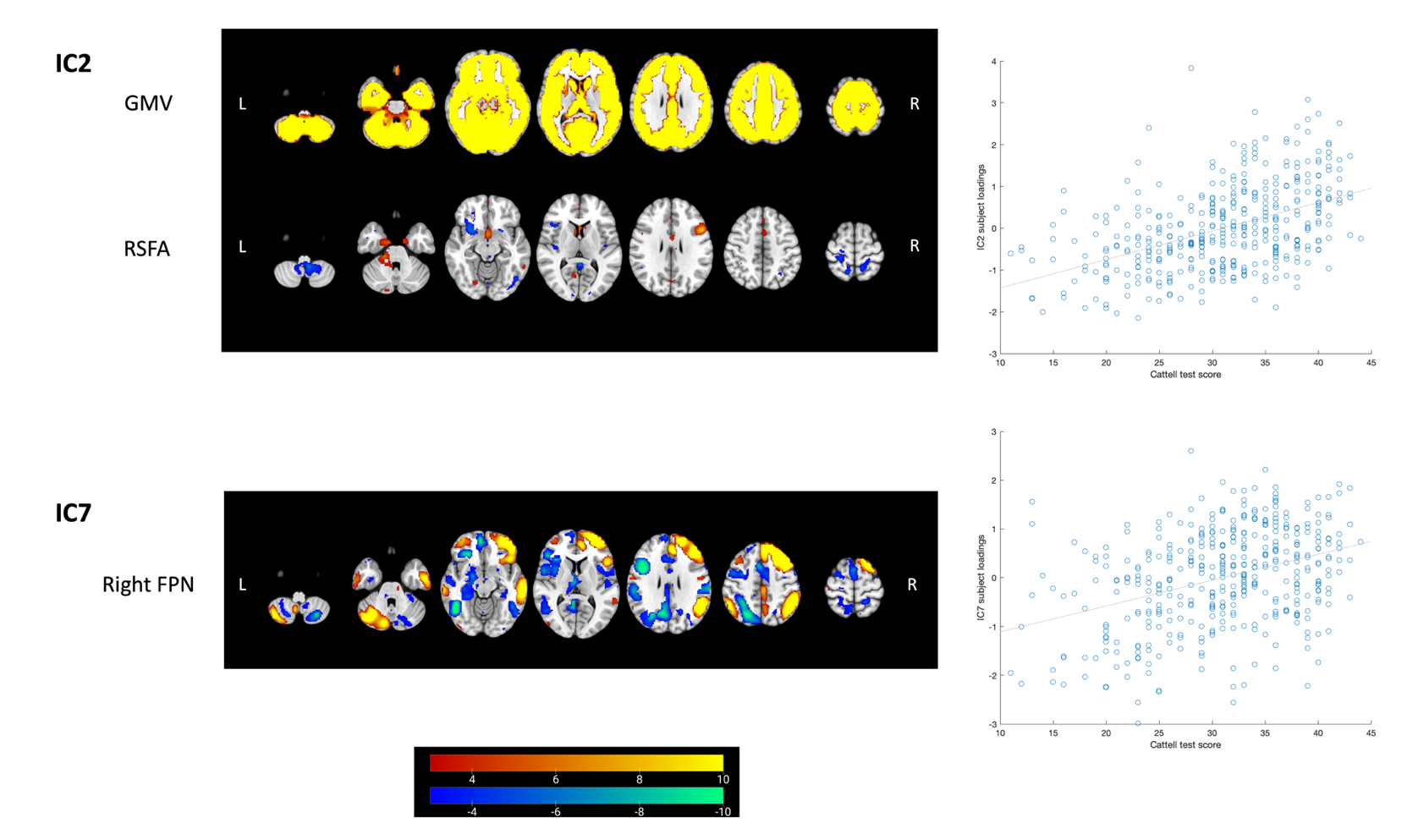

**Supplementary Figure 2.** Linked ICA weighted spatial maps for two components showing unique associations with Cattell test score in CC420 analysis (*n* = 433). Warm and cold colour scheme indicate positive and negative association with linked ICA subject loadings, respectively. The accompanying scatter plots show models of Cattell test score plotted against IC subject loadings. One component reflects signals from structural and cerebrovascular measurements: IC2 which reflects the grey matter volume (GMV) and resting state fluctuation amplitude (RSFA). One component, IC7 which reflects the right frontoparietal network (FPN), showed an interaction between Cattell test score and age in multiple regression analysis (supplementary table). Two similar components were found in CC280 main analysis to be associated with fluid intelligence. For visualization the spatial map threshold is set to 3 < |*Z*| < 10.

**Supplementary Table.** Multiple regression analysis results of each independent component (IC) subject loadings from linked independent component analysis of CC420 participants (40 components based on 6 modalities and *n* = 433).

|  | IC ~ Cattell*Age + Age^2 + gender + head motion | | | | | | | | | | |
| --- | --- | --- | --- | --- | --- | --- | --- | --- | --- | --- | --- |
| IC | Overall model fit | | | Age | | Age^2 | | Cattell | | Cattell*Age | |
|  | Adjusted R^2^ | *P* | FWER-corrected *P* | t | *P* | t | *P* | t | *P* | t | *P* |
| IC1 | 0.16 | < 0.0001 | **< 0.0001** | -5.41 | **< 0.0001** | 2.19 | **0.029** | 0.59 | 0.56 | -0.69 | 0.49 |
| IC2 | 0.44 | < 0.0001 | **< 0.0001** | -6.30 | **< 0.0001** | 1.74 | 0.083 | 4.26 | **< 0.0001** | -0.73 | 0.47 |
| IC3 | 0.039 | 0.0011 | **0.044** | -2.69 | **0.0075** | -0.54 | 0.59 | 0.78 | 0.43 | -0.14 | 0.89 |
| IC4 | 0.0028 | 0.31 | > 0.99 |  |  |  |  |  |  |  |  |
| IC5 | 0.27 | < 0.0001 | **< 0.0001** | 3.38 | **0.00080** | -0.045 | 0.96 | -1.68 | 0.094 | 0.080 | 0.94 |
| IC6 | 0.054 | < 0.0001 | **0.0026** | -1.16 | 0.25 | 1.49 | 0.14 | -0.13 | 0.90 | 1.37 | 0.17 |
| IC7 | 0.28 | < 0.0001 | **< 0.0001** | -3.02 | **0.0027** | 0.68 | 0.50 | 1.37 | 0.17 | 2.98 | **0.0031** |
| IC8 | -0.0041 | 0.64 | > 0.99 |  |  |  |  |  |  |  |  |
| IC9 | 0.73 | < 0.0001 | **< 0.0001** | -21.49 | **< 0.0001** | -8.25 | **< 0.0001** | -0.15 | 0.88 | -0.59 | 0.56 |
| IC10 | 0.27 | < 0.0001 | **< 0.0001** | -4.01 | **< 0.0001** | 0.17 | 0.86 | 1.16 | 0.25 | 1.88 | 0.061 |
| IC11 | 0.19 | < 0.0001 | **< 0.0001** | -2.70 | **0.0073** | -0.042 | 0.97 | 0.00087 | > 0.99 | 1.87 | 0.062 |
| IC12 | 0.0084 | 0.15 | > 0.99 |  |  |  |  |  |  |  |  |
| IC13 | 0.0057 | 0.22 | > 0.99 |  |  |  |  |  |  |  |  |
| IC14 | 0.22 | < 0.0001 | **< 0.0001** | -2.51 | **0.012** | 0.84 | 0.40 | -1.49 | 0.14 | -0.27 | 0.79 |
| IC15 | 0.017 | 0.041 | > 0.99 |  |  |  |  |  |  |  |  |
| IC16 | 0.017 | 0.044 | > 0.99 |  |  |  |  |  |  |  |  |
| IC17 | 0.067 | < 0.0001 | **0.00023** | -3.15 | **0.0017** | 1.06 | 0.29 | 0.28 | 0.78 | -0.084 | 0.93 |
| IC18 | 0.046 | 0.00033 | **0.013** | -0.82 | 0.41 | -3.22 | **0.0014** | 0.87 | 0.38 | -1.33 | 0.18 |
| IC19 | 0.14 | < 0.0001 | **< 0.0001** | -0.68 | 0.50 | -2.27 | **0.024** | 0.85 | 0.40 | -0.26 | 0.79 |
| IC20 | 0.039 | 0.0010 | **0.040** | -0.99 | 0.32 | -0.49 | 0.62 | -1.79 | 0.075 | -0.38 | 0.70 |
| IC21 | 0.076 | < 0.0001 | **< 0.0001** | 1.84 | 0.066 | -2.35 | **0.019** | -0.68 | 0.50 | 0.90 | 0.37 |
| IC22 | 0.049 | 0.00018 | **0.0072** | 1.68 | 0.094 | -0.85 | 0.40 | -1.64 | 0.10 | 0.28 | 0.78 |
| IC23 | 0.12 | < 0.0001 | **< 0.0001** | -4.93 | **< 0.0001** | -2.38 | **0.018** | -0.52 | 0.61 | -1.71 | 0.087 |
| IC24 | < 0.0001 | 0.43 | > 0.99 |  |  |  |  |  |  |  |  |
| IC25 | 0.27 | < 0.0001 | **< 0.0001** | -2.61 | **0.0093** | -10.46 | **< 0.0001** | -0.93 | 0.35 | -1.11 | 0.27 |
| IC26 | 0.10 | < 0.0001 | **< 0.0001** | 3.49 | **0.00054** | -0.97 | 0.33 | -1.06 | 0.29 | 1.40 | 0.16 |
| IC27 | 0.014 | 0.066 | > 0.99 |  |  |  |  |  |  |  |  |
| IC28 | 0.035 | 0.0023 | 0.092 |  |  |  |  |  |  |  |  |
| IC29 | 0.011 | 0.10 | > 0.99 |  |  |  |  |  |  |  |  |
| IC30 | 0.044 | 0.00047 | **0.019** | -0.42 | 0.67 | 0.73 | 0.46 | -1.10 | 0.27 | 0.42 | 0.67 |
| IC31 | 0.0067 | 0.19 | > 0.99 |  |  |  |  |  |  |  |  |
| IC32 | 0.014 | 0.071 | > 0.99 |  |  |  |  |  |  |  |  |
| IC33 | 0.0016 | 0.36 | > 0.99 |  |  |  |  |  |  |  |  |
| IC34 | 0.13 | < 0.0001 | **< 0.0001** | 2.42 | **0.016** | -1.34 | 0.18 | 0.30 | 0.76 | -1.89 | 0.060 |
| IC35 | 0.0074 | 0.17 | > 0.99 |  |  |  |  |  |  |  |  |
| IC36 | 0.0077 | 0.17 | > 0.99 |  |  |  |  |  |  |  |  |
| IC37 | 0.0037 | 0.28 | > 0.99 |  |  |  |  |  |  |  |  |
| IC38 | 0.00013 | 0.42 | > 0.99 |  |  |  |  |  |  |  |  |
| IC39 | -0.0080 | 0.85 | > 0.99 |  |  |  |  |  |  |  |  |
| IC40 | -0.0073 | 0.81 | > 0.99 |  |  |  |  |  |  |  |  |

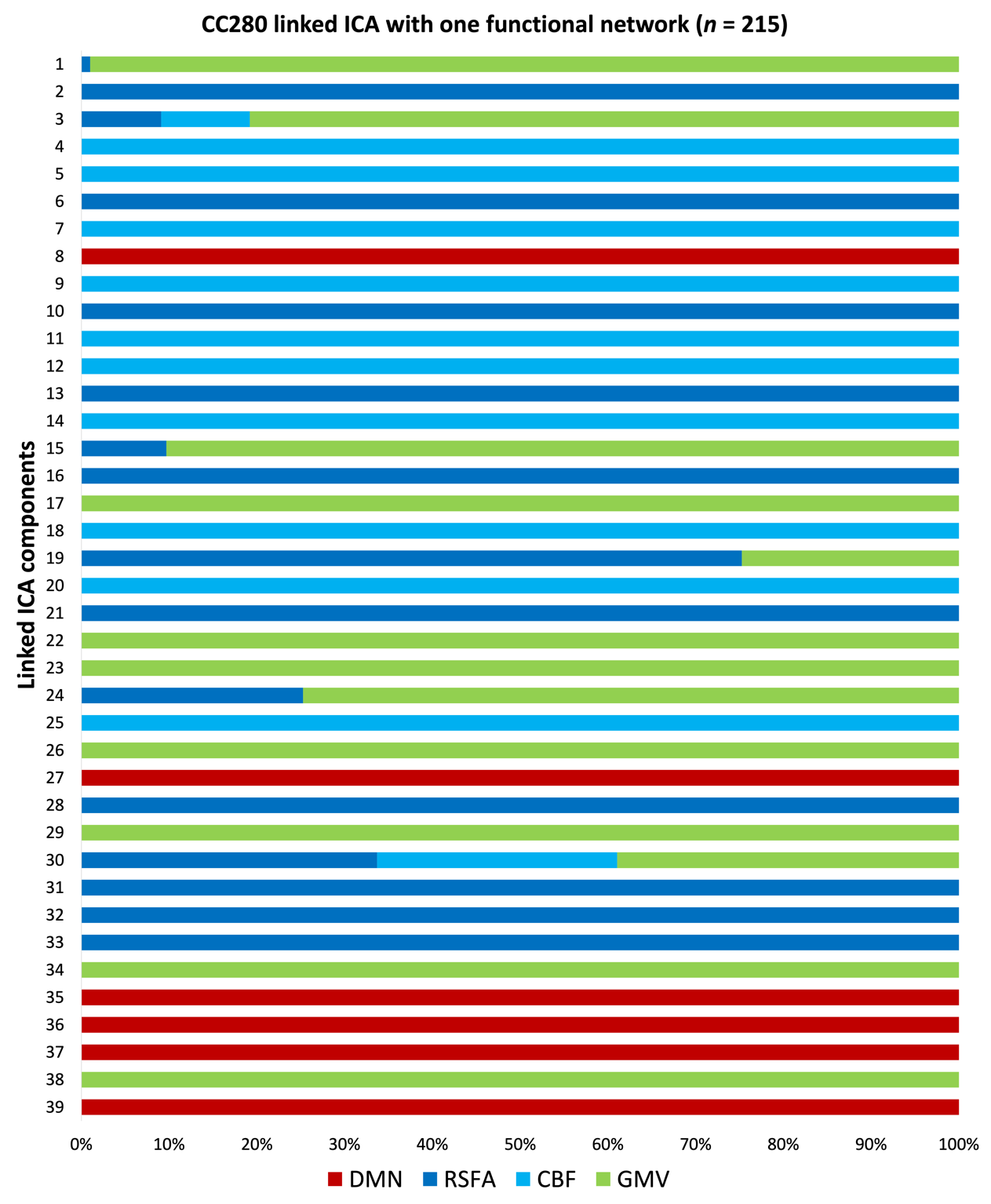

**Supplementary Figure 3.** The relative weight of modalities in each component generated from linked independent component analysis (ICA) with one functional network (*n* = 215). No fusion was found between the DMN and other modalities. Abbreviations: DMN, default mode network; RSFA, resting state fluctuation amplitude; CBF, cerebral blood flow; GMV, grey matter volume.
